# Sub-threshold neuronal activity and the dynamical regime of cerebral cortex

**DOI:** 10.1101/2022.07.14.500004

**Authors:** Oren Amsalem, Hidehiko Inagaki, Jianing Yu, Karel Svoboda, Ran Darshan

## Abstract

Cortical neurons exhibit temporally irregular spiking patterns and heterogeneous firing rates. These features arise in model circuits operating in a ‘fluctuation-driven regime’, in which fluctuations in membrane potentials emerge from the network dynamics. However, it is still unclear whether the cortex operates in this regime. We evaluated the fluctuation-driven hypothesis by analyzing spiking and sub-threshold membrane potentials of neurons in the sensory and frontal cortex recorded during a decision-making task. Standard fluctuation-driven models account for spiking statistics but fail to capture the heterogeneity in sub-threshold activity. We address this issue by effectively incorporating dendritic conductances into the standard models. Our model suggests that the frontal cortex operates in a fluctuation-driven regime. In contrast, excitatory neurons in layer 4 of the barrel cortex are not fluctuation-driven; they spike in response to occasional synchronous inputs. Our work reveals fundamental differences between cortical areas, suggesting that they operate in different dynamical regimes.

## Introduction

The cortex operates in a noisy dynamical regime. Cortical neurons spike at irregular times, with statistics that can be approximated by a Poisson process [Softky and Koch, 1993, Shadlen and Newsome, 1998, Compte et al., 2003]. In addition, neuronal firing rates are highly heterogeneous, with some that fire at high rates while a large number of neurons that are almost quiescent [Griffith and Horn, 1966, Hromádka et al., 2008, O’Connor et al., 2010, Buzsáki and Mizuseki, 2014]. Ongoing research in system neuroscience is directed at understanding this operating regime of cortex [Ahmadian and Miller, 2021], in which spiking irregularity and rate heterogeneity are ubiquitous.

Sensory input and movement can contribute to irregular spiking and heterogeneous spike rates [Hires et al., 2015, Stringer et al., 2019]. However, substantial contribution to the variability and heterogeneity likely results from the recurrent network dynamics. Indeed, theoretical studies have shown that these features can emerge from the non-linear recurrent dynamics of cortical networks, if they operate in a ‘fluctuation-driven regime’ in which excitatory and inhibitory currents are both large compared to the rheobase of the neurons but approximately balance each other [van Vreeswijk and Sompolinsky, 1996, Amit and Brunel, 1997, Brunel, 2000, Hertz et al., 2003]. The net input can be sub-threshold, on average, while action potentials are driven by fluctuations in synaptic inputs [Vogels et al., 2005].

Theoretical studies of the fluctuation-driven regime are foundational to our quantitative understanding of spiking statistics in the cortex [Van Vreeswijk and Sompolinsky, 2005, Renart et al., 2010, Roxin et al., 2011, Hansel and Mato, 2013, Rosenbaum et al., 2017, Darshan et al., 2018]. However, it is still debated whether the cortex operates in this regime [Ahmadian and Miller, 2021]. For example, fluctuation-driven networks can quantitatively account for feature selectivity in the cortex [Hansel and van Vreeswijk, 2012, Pehlevan and Sompolinsky, 2014, Pattadkal et al., 2018] and are consistent with cortical responses to perturbation experiments in sensory and frontal areas [Mahrach et al., 2020, Sanzeni et al., 2020, Kim et al., 2022]. On the other hand, similar perturbation experiments in layer 4 neurons of the barrel cortex are inconsistent with a fluctuation-driven hypothesis [Gutnisky et al., 2017]. Furthermore, while some modeling studies suggest that recurrent and external inputs to cortical neurons are large [Hansel and van Vreeswijk, 2012, Mahrach et al., 2020, Sanzeni et al., 2020], it has been argued that feedforward thalamic inputs to the visual cortex are too weak to be approximately balanced by a strong recurrent inhibition [Ahmadian and Miller, 2021].

Classical models of networks that operate in the fluctuation-driven regime, in which synaptic interactions are mediated by variations in the synaptic conductance, predict that neuronal membrane potential should hover close to the neuronal threshold [Tan et al., 2014] and, as we will show, with limited heterogeneity in mean voltage across neurons. In this study, we tested the fluctuation-driven regime hypothesis by analyzing the *supra-* and *sub-threshold* activity of populations of excitatory and inhibitory neurons in the anterior lateral motor cortex (ALM) and in the vibrissal somatosensory area (vS1, or barrel cortex) in behaving mice that perform a decision-making task. Does the same mechanism for explaining spiking statistics in cortex accounts for the variability and heterogeneous sub-threshold voltage activity?

We show that fluctuation-driven networks can account for spiking statistics in ALM. However, they fail to reproduce the large level of voltage heterogeneity observed in the data due to the strong excitatory and inhibitory synaptic inputs that induce a large increase in the neuronal input conductance. We resolve this discrepancy by introducing a phenomenological network model of *‘extended-like’* point neurons with synapses that mimic dendritic integration, thus including a key aspect of neuronal biophysics that is often neglected in models of recurrent neural networks [Larkum, 2022]. This model suggests that neurons in ALM are fluctuation-driven. This is also the case for the inhibitory neurons in the barrel cortex. In contrast, our modeling and analysis show that excitatory neurons in layer 4 of the barrel cortex are not fluctuation-driven. Their mean voltage is far from the threshold and they spike in response to occasional synchronous input. We therefore suggest that layer 4 neurons in the barrel cortex are only *partially-balanced*, meaning that the excitatory neurons are unbalanced and are dominated by inhibition, while the thalamic excitatory and recurrent inhibitory currents to the inhibitory neurons are approximately balanced. Our work suggests that during decision making cortical excitatory neurons closer to the periphery are not balanced and they fire due to strong and correlated external drive, whereas neurons in other populations hover closer to their thresholds, their currents are approximately balanced, and they operate in a fluctuation-driven regime.

## Results

### Spiking and sub-threshold activity in ALM

Mice were engaged in a sensorimotor task: they responded to an instruction cue, presented during a sample epoch, by licking for a water reward following a second-long delayed epoch (Fig.1, [Guo et al., 2014]). We analyzed the supra (spiking) and sub-threshold activity (membrane potential) of individual excitatory and inhibitory cells [Guo et al., 2017, Inagaki et al., 2019]. In ALM a large proportion of recorded neurons exhibited preparatory activity that predicts licking direction [Inagaki et al., 2019].

**Figure 1:**
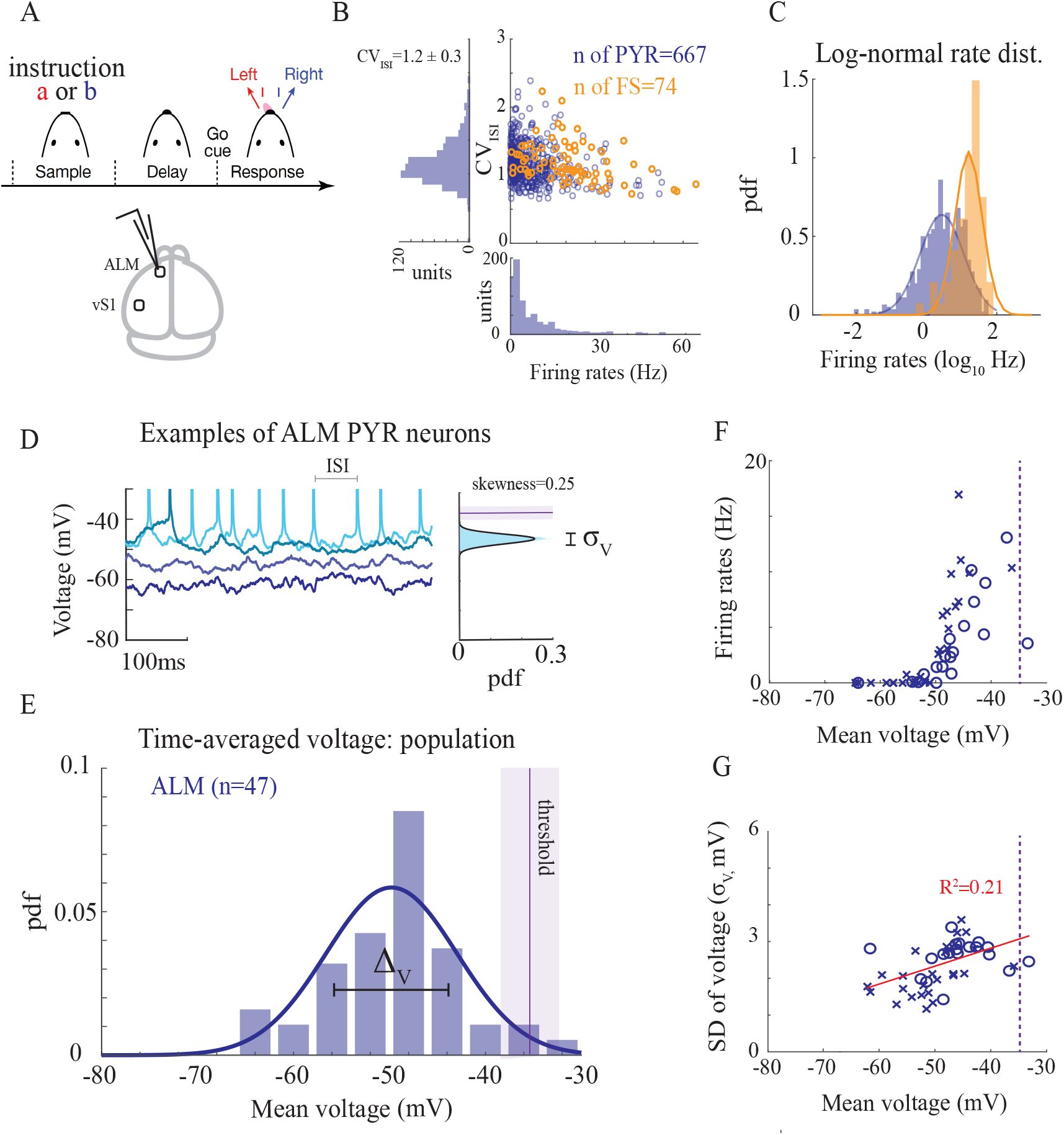
Supra- and sub-threshold statistics of ALM pyramidal neurons in mice performing a delayed-response task. **A**. Top: behavioral task. Bottom: recording area. **B**. *CV*_*ISI*_ against neuronal firing rates for putative pyramidal (PYR) and fast-spiking (FS) inhibitory neurons (FS). **C**. Probability density function (pdf) of the log-rates is well-approximated by a Gaussian distribution (solid lines). **D**. Left: Activity of four example neurons during the first 0.5 second of delay period, together with an illustration of the inter-spike-interval. Right: Sub-threshold voltage distribution (excluding spikes) of one of the neurons. Solid line: fit to a Gaussian distribution. Purple line and shaded areas: neuronal threshold (mean± SD). *σ*_*V*_ : SD of the single-neuron fluctuations. **E**. pdf of mean voltage of the *n* = 47 recorded ALM neurons. Solid line: fit to a Gaussian distribution. Purple line and dashed area around it: average threshold across the population and its SD (−34 ± 3*mV*) **F**. Firing rates vs. mean voltage. Crosses: auditory instruction (28 neurons). Circles: tactile instruction (19 neurons) (see Methods). **G**. SD of single-neuron voltage fluctuations against the mean voltage. Red line: linear regression. In all panels the data is analyzed for correct lick-right trials during delay period. In (B-C) Extracellular recordings (silicon probes). In (D-G) whole-cell recordings.

Consistent with previous recordings in other brain areas and species [Shinomoto et al., 2003, Buzsáki and Mizuseki, 2014], we found that spiking statistics of putative-pyramidal (PYR) and fast-spiking (FS) neurons exhibited large temporal irregularities, high trial-to-trial variability, heterogeneous spike rates and diverse selective responses. Distributions of firing rates of pyramidal neurons were approximately log-normal (Fig.1B-C). Spiking activity exhibited wide inter-spike-interval (ISI) distributions, with average coefficient of variations of the ISIs that varied between behavioral states (Fig.1B, *CV*_*ISI*_ = 0.9 during sample, and *CV*_*ISI*_ = 1.2 during delay; paired Student t-test *p <* 0.001), but were close to one (for comparison, neurons that fire regularly have low *CV*_*ISI*_, while for a Poisson neuron *CV*_*ISI*_ = 1). Importantly, the *CV*_*ISI*_ was large for neurons of both low and high spike rates. FS neurons fired, on average, at higher rates than pyramidal neurons and their rate distribution was also well-approximated by a log-normal distribution. Similarly, their spiking activity was highly variable, with *CV*_*ISI*_ that was slightly higher than the *CV*_*ISI*_ of the pyramidal neurons. Neurons in both populations exhibited a high level of variability across trials (Fig.S5B). This activity was selective to licking direction and was diverse across the population ([Inagaki et al., 2019]; Fig.S5B).

We next examined membrane potential measurements from whole-cell recordings of ALM neurons (Fig.1D-G; [Guo et al., 2017, Inagaki et al., 2019]). The sub-threshold fluctuations of most of the recorded neurons were well-approximated by a Gaussian distribution (excluding their spikes, example in Fig.1D; skewness of distribution was 0.5 ± 0.44, mean± SD; Fig.S2; [Tan et al., 2014]). The time-average voltage across the population was −50*mV* (range: −64.5*mV* to −33.5*mV*). There was a large level of heterogeneity around this average value (Fig.1E-G), with only a small fraction of the neurons that were close to their threshold (threshold was −34 ± 3*mV*). We quantified this heterogeneity in sub-threshold statistics by the standard deviation of the time-average voltages across the population, Δ_*V*_, which was around 6*mV* for ALM neurons. For neurons that spiked (*n* = 35*/*47 neurons), when compared to their thresholds, the mean voltage ranged from 6*mV* to 25*mV* below threshold (with mean of 13*mV* and SD of 4*mV*).

The level of fluctuations of sub-threshold activity of neurons around their temporal-averaged value varied across the population (standard deviation of voltage fluctuations, *σ*_*V*_, is in the range 1.2 − 4.5*mV*). Neurons close to threshold exhibited a larger level of voltage fluctuations. The level of fluctuations and mean voltage were positively correlated across the population (Fig.1G). Finally, the mean voltage of neurons was selective to licking direction. Similarly to the supra-threshold selectivity, such selectivity at the level of the average voltage was diverse as well (Fig.S5B-C).

### Spiking and sub-threshold activity in vS1

We then analyzed the supra- and sub-threshold activity of neurons in the vibrissal somatosensory area (vS1) during the sampling period in a similar decision making task (a Go/NoGo task lacking a delay period, Fig.2A, [Yu et al., 2016]). We excluded periods of touch, as the activity of neurons in vS1 varied substantially when a whisker touched an object.

**Figure 2:**
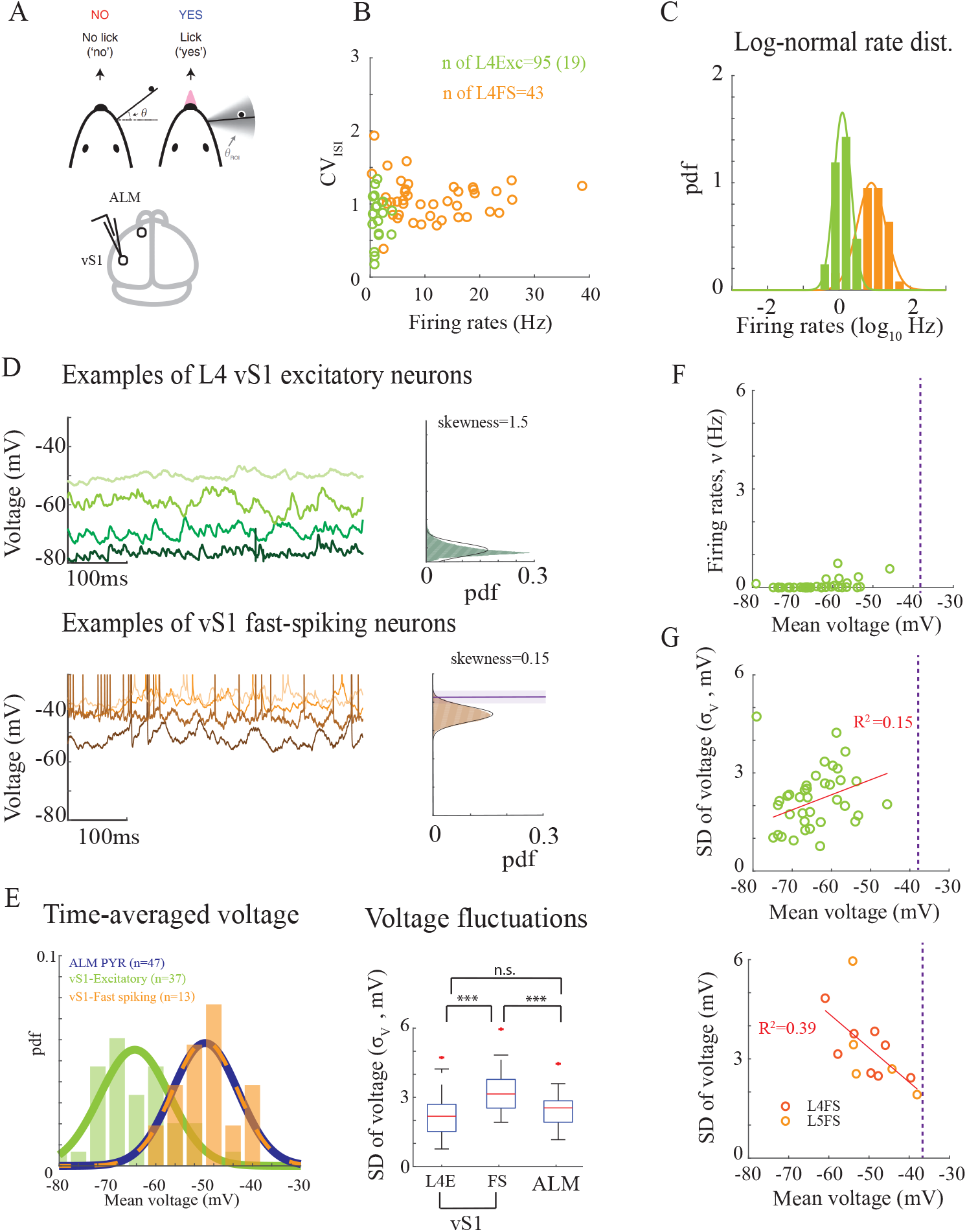
Supra- and sub-threshold statistics of vS1 neurons in mice performing a Go/NoGo task. **A**. Top: behavioral task. Bottom: recording area. **B**. *CV*_*ISI*_ against neuronal firing rates for neurons in layer 4. Most of the excitatory neurons were quiescent (only 19/95 neurons that spiked are presented). **C**. Probability density function (pdf) of the log-rates. Solid lines: fit to a Gaussian distribution. **D**. Top: Examples of layer 4 excitatory neurons. Bottom: Fast-spiking neurons. Left: Activity of example neurons during the first half-second of a whisking episode. Right: Sub-threshold voltage distribution of one of the neurons. Solid line: fit to a Gaussian distribution. **E**. Left: Probability density function (pdf) of time-average voltage for all recorded neurons. Solid lines: fit to a Gaussian distribution. Right: SD of (single neuron) voltage fluctuations for all recorded neurons. **F**. Firing rates vs. mean voltage for the excitatory neurons. **G**. SD of (single neuron) voltage fluctuations against the mean voltage. Top: layer 4 excitatory neurons. Bottom: Fast-spiking neurons in layer 4 and Layer 5. Red line: linear regression. In (B-C) Extracellular and intracellular recordings. In (D-G) whole-cell recordings. In (B-C,E-G) analysis during non-whisking periods.

Similar to neurons in ALM, neurons in vS1 fired with high *CV*_*ISI*_. Their firing rate statistics were well-approximated by a log-normal distribution, and FS neurons fired at higher rates than the excitatory neurons (Fig.2B-D, Fig.S1B-D). When considering the sub-threshold activity of the excitatory population of vS1, we found that their resting state was much more hyperpolarized than the excitatory neurons in ALM (Fig.2D top, Fig.2E; Student t-test, *p <* 0.01). Separating the excitatory neurons in vS1 to layer 4 (L4E) and layer 5 (L5E) neurons, we found that L4E neurons were much more hyperpolarized than ALM and L5E neurons (Fig.2E, Fig.S1E). The distance to threshold of L4E neurons was 21*mV* ± 7*mV* (for *n* = 19*/*37 cells that fired action potentials; threshold was −39*mV* ± 5*mV*) and 17*mV* ± 7*mV* for L5E neurons (for *n* = 29*/*38 cells that fired action potentials; threshold was −35 ± 5*mV*). As a result of their large distance to threshold, L4E neurons spiked at extremely low rates and almost exclusively as a result of touch ([Yu et al., 2016]; Fig.2B). In contrast to ALM neurons, distributions of voltage fluctuations of the excitatory neurons in vS1 tended to deviate from a Gaussian distribution (example of distribution in Fig.2D; average skewness of distribution was 1.0 ± 0.6 for L4E neurons and 0.9 ± 0.6 for L5E neurons). Skewness of the distribution and resting potential, especially for L4E neurons, were anti-correlated (Fig.S2B), with positively skewed voltage distributions of neurons that were farther from threshold.

Besides the differences in the resting state, sub-threshold activity of excitatory neurons in ALM and vS1 also shared common features. First, as with ALM neurons, there was a large level of heterogeneity in the mean voltage of the neurons (large Δ_*V*_), with some neurons much closer to their threshold than others. Second, we found that both in ALM and vS1 the level of fluctuations was quite similar (Fig.2E, Student t-test, *p >* 0.05) and their voltage SDs and means were positively correlated across the population (Fig.2G-top, Fig.S1G).

In vS1 we were also able to record the sub-threshold activity of 13 FS neurons (combining eight layer 4 and five layer 5 FS neurons). We compared their activity to the sub-threshold activity of excitatory neurons in vS1. FS neurons were more depolarized than the excitatory neurons, and their voltage fluctuations were larger, which contributed to their higher spike rate. When compared with ALM neurons, we found that the mean voltages of FS neurons in vS1 were not significantly different than the mean voltages of pyramidal neurons in ALM (Fig.2E-left, distance to threshold: 12*mV* ± 5*mV*, Student t-test, *p >* 0.05, threshold −37*mV* ± 4 mV) and the distribution of their voltage fluctuations were also well-fitted by a Gaussian distribution (skewness of 0.3 ± 0.3). However, FS neurons exhibited a larger level of voltage fluctuations (Fig.2E-right) than ALM. Finally, in contrast to the excitatory populations, FS neurons exhibited negative correlations between their SDs and mean voltages (Fig.2G-bottom).

### Spiking network models that operate in a fluctuation-driven regime are inconsistent with the sub-threshold statistics of cortical neurons

We next asked if network models that operate in the fluctuation-driven regime could account for the spiking as well as the sub-threshold activity of neurons in the data. To answer this question, we considered a network consisting of excitatory and inhibitory integrate-and-fire neurons, where cells were randomly and sparsely connected to each other (Fig.3). The post-synaptic neuronal voltage, *V* (*t*), evolved in time and changed in response to synaptic currents. These currents were modeled as *I*_*syn*_(*t*) = −*gs*(*t*)(*V* (*t*)−*E*), with *E* being the synaptic reversal potential, *g* was the change in synaptic conductance induced by a pre-synaptic spike, and *s*(*t*) was a filtered version of the pre-synaptic spikes (see Methods for a complete description of the model). These type of network models are known in the literature as *conductance-based networks* because variations in synaptic inputs lead to changes in the conductance of the neuron [Destexhe et al., 2003, Hertz et al., 2003, Richardson, 2004, Sanzeni et al., 2022].

**Figure 3:**
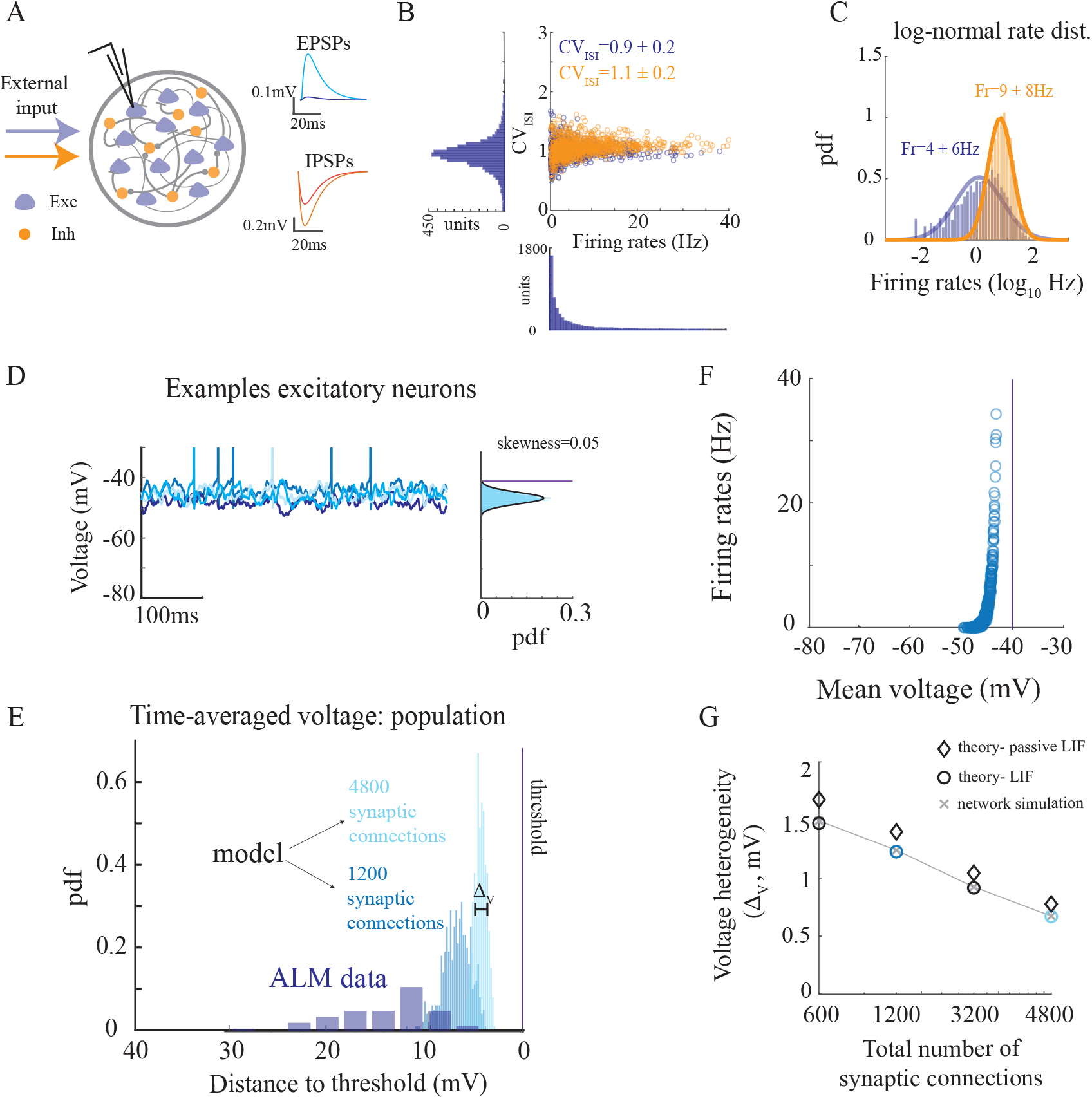
A network of one-compartment integrate-and-fire neurons is consistent with irregular spiking statistics in ALM but fails to reproduce sub-threshold voltage statistics. **A**. Left: Diagram of the recurrent neural network. Synapses in the network are mediated by variations in the synaptic conductances. Right: excitatory and inhibitory post-synaptic potentials in the network. Cyan: Inh-to-Exc connection. Blue: Exc-to-Exc. Red: Exc-to-Inh. Orange: Inh-to-Inh. **B**. *CV*_*ISI*_ vs. firing rate for excitatory (blue) and inhibitory (orange) neurons in the network. **C**. Distributions of firing rates in the network. **D**. Left: Activity of four example neurons. Right: Sub-threshold voltage distribution (excluding spikes) of one of the neurons. Solid line: fit to a Gaussian distribution. Purple line: neuronal threshold. **E**. Distribution of time-average voltage, plotted as the distance to threshold (purple line) of all neurons in a network with an average of 1200 (blue) and 4800 (cyan) pre-synaptic inputs per neuron (total synaptic connections: *K*_*I*_ + 2*K*_*E*_ ; Methods). Threshold in simulations was at −40*mV*. ALM data from Figure Fig.1E is re-plotted here as the distance to neuronal threshold for comparison. **F**. Firing rate vs. mean voltage for all neurons in the network of a total 1200 pre-synaptic inputs. **G**. Voltage heterogeneity vs. number of average pre-synaptic inputs. Crosses: network simulations. Diamonds: predictions using of a passive neuron (first term in Eq.(1)). Circles: predictions using an integrate-and-fire neuron, including threshold (full expression in Eq.(1)).

The distributions of firing rates for both excitatory and inhibitory populations were well approximated by a log-normal distribution (Fig.3B-C; [Roxin et al., 2011, Sanzeni et al., 2022]). We chose the synaptic parameters such that the distributions of neuronal firing rates in the populations fit well the spiking data of ALM neurons. In particular, the firing rate of the inhibitory population was larger than the excitatory one. With these parameters, the size of the excitatory and inhibitory post-synaptic potentials were within the physiological range (Fig.3A). Neurons in both populations fired irregularly and with *CV*_*ISI*_ ≈ 1, as shown in the data. It was slightly higher for the inhibitory neurons than the excitatory one. This variability was self-generated by the network. The fact that the *CV*_*ISI*_ was large for both neurons that fired at low and high rates is a hallmark of the fluctuation-driven regime, where fluctuations are driving the spikes [van Vreeswijk and Sompolinsky, 1996].

We next investigated the sub-threshold statistics of the simulated neurons. In contrast to the spiking statistics of neurons in the network, we found that the network model failed to reproduce the sub-threshold statistics of ALM neurons. Specifically, the time-average voltage of all neurons in the model was close to their threshold, with very small standard deviation (SD) around the average value (Fig.3D-E; mean voltage of −45*mV* with threshold at −40*mV* and SD of Δ_*V*_ = 1*mV*). Similarly, the amplitude of the temporal voltage fluctuations across the population was narrowly distributed (*σ*_*V*_ ≈ 2*mV* for all neurons). This is in contrast to the large heterogeneity in mean voltages and SDs across the ALM neurons (Fig.3E-F, Fig.1E,G).

How is it that firing rates in the simulated networks were so heterogeneous while the time-average voltage distribution was so narrow? We found that this effect was a direct outcome of the high-conductance regime in which the network operated. Indeed, opening of a synaptic channel increases the effective conductance– the ‘leak’, of the neuron. Due to a large number of open synaptic channels, the neuron becomes very leaky and its time-average voltage is clamped to an effective reversal. This effective reversal depends on the time-average conductances and reversal potentials of its excitatory and inhibitory synapses. Moreover, with voltage fluctuations of ∼ 2*mV*, this time-average voltage must be very close to threshold in order for the neuron to fire. The fact that the firing rates are highly heterogeneous is remarkable, and it results from the large gain of the firing-voltage curve in these models ([Richardson, 2004], Fig.3F).

To make this argument quantitative, we studied analytically the heterogeneity in the sub-threshold activity of one-compartment leaky integrate-and-fire neurons. We used a mean-field theory to determine the response of a population of non-interacting neurons to inputs from populations of excitatory and inhibitory neurons ([Richardson, 2004, Sanzeni et al., 2022], Methods and Supplementary materials). Pre-synaptic inputs were modeled as Poisson processes, with heterogeneous rate distribution (e.g., log-normal distribution). We determined the distribution of the voltage of the non-interacting neurons as a function of the number, strength and rate distribution of their pre-synaptic inputs. Using this approach, the time-average voltage of a post-synaptic neuron, 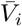, is (see Methods):

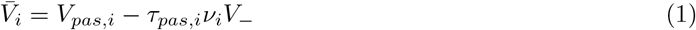

with *ν*_*i*_ being the firing rate of the *i*’th neuron and *V*_*−*_ = *V*_*th*_ − *V*_*r*_ is the difference between the threshold (*V*_*th*_) and the resetting voltage (*V*_*r*_). The level of sub-threshold heterogeneity is thus a result of two terms; the first term in Eq.(1), *V*_*pasi,i*_, is a result of bombarding a passive neuron with a large number of synapses that ‘shunts’ the soma and clamps the neurons to the same resting state. In large networks it is not very different across neurons (Methods). The second term of Eq.(1) depends on the neuronal firing rate, which varies considerably across neurons in the network. Each spike results in resetting the voltage from the neuronal threshold to its reset value, contributing to the voltage a factor of *τ*_*pas,i*_*V*_*−*_, with *τ*_*pas,i*_ being the neuronal effective timescale. This timescale depends on the network state and it is small when the number of pre-synaptic inputs to the neuron is large (see Methods). Taken together, the distribution of mean voltages across the population in such networks is expected to be very narrow. It vanishes with increasing number of synaptic inputs (Figure 3E,G).

To conclude, classic network models that operate in the fluctuation-driven regime were consistent with spiking statistics in the cortex, but fundamentally failed to account for heterogeneity in sub-threshold statistics. The low level of voltage heterogeneity across neurons was an inevitable result of the high-conductance regime in which the network operated that clamped the voltage of all the neurons to more or less the same effective reversal.

### A phenomenological point neuron model that mimics an effective dendritic morphology

In the previous section we considered a recurrent network consisting of neurons modeled as single-compartment elements (point neurons) [Destexhe et al., 2003, Richardson, 2004, Sanzeni et al., 2022]. Because of this simplification, all activated synapses contributed the same amount of change to the neuronal conductance. When many synapses were active at the same time, all neurons in the network were clamped to more or less the same value of membrane potential. However, in reality, neurons exhibit extended spatial structures and the effect of synaptic inputs on somatic membrane conductance depends on their proximity to the soma [Rall, 1964, Koch et al., 1990, Destexhe et al., 2003]. We hypothesized that the shunting effect that led to the narrow voltage distribution in our networks would be less pronounced if the neuronal model took into account the neuronal morphology.

Most of the putative pyramidal cells we recorded in ALM were layer 5 neurons. Thus, to test our hypothesis, we first simulated a reconstructed layer 5 pyramidal neuron [Scala et al., 2021, Ascoli et al., 2007] (Fig.4A-B), in which the spiking activities of its pre-synaptic neurons were Poisson. For each pre-synaptic excitatory (inhibitory) neuron we sampled its spike rate from the log-normal distribution of ALM pyramidal (fast-spiking) neurons (see Fig.1B,C). We repeated this process 5000 times, each time sampling a set of pre-synaptic neurons with different spike rates from the log-normal rate distribution, and used it to calculate the distribution of mean voltages over the population of 5000 neurons. When localizing all synapses on the soma (Fig.4A) we found that, similarly to what happens in a network of point neurons, the distribution of mean voltages across the 5000 realizations of the pyramidal neuron was extremely narrow (Fig.4C). Voltage fluctuations (*σ*_*V*_), as well as voltage heterogeneity (Δ_*V*_) both decreased with increasing number of synaptic inputs (gray lines in Fig.4D). However, when distributing the same number of synapses on the dendritic tree (Fig.4B), we observed that the width of the time-averaged voltage distribution increased significantly (Fig.4C). Both the voltage heterogeneity and the size of the fluctuations weakly depended on the number of synaptic inputs (black lines Fig.4D). These results suggest that voltage heterogeneity can be significant in large model networks that operate in a fluctuation-driven regime, provided that one takes into account the neuronal morphology.

**Figure 4:**
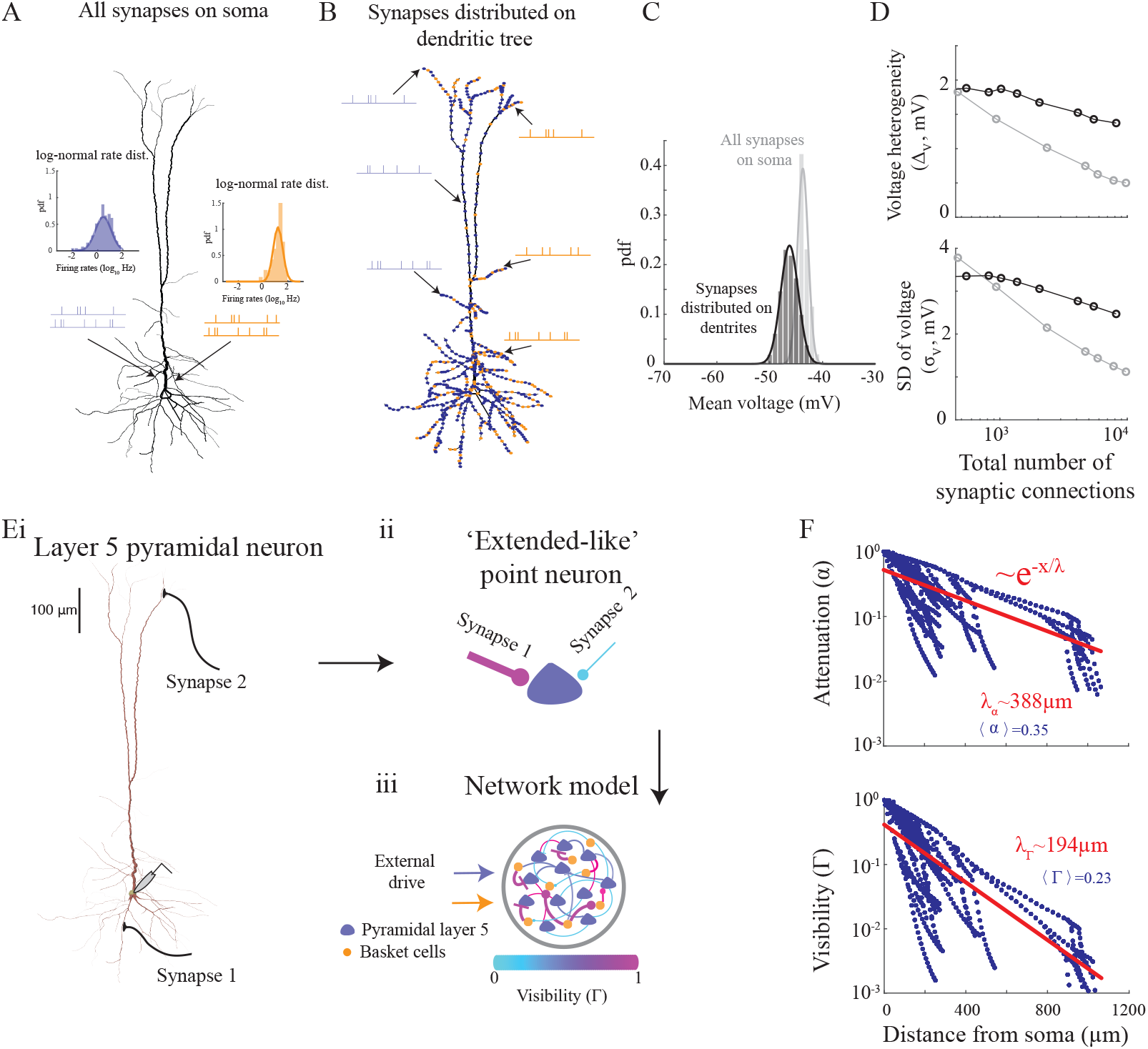
Sub-threshold heterogeneity and the effect of distribution of synaptic inputs in a multi-compartment model of layer 5 pyramidal neuron. **A**. Simulations of a layer 5 pyramidal neuron with all AMPA (blue) and GABAA (orange) synapses located on the soma. Pre-synaptic Poisson neurons are simulated with firing rates sampled from the log-normal rate distribution of the pyramidal (blue) and fast-spiking (orange) ALM neurons. 75% excitatory and 25% inhibitory pre-synaptic neurons are simulated for each cell. **B**. Same as (A) but when synapses are uniformly distributed along the dendritic tree. **C**. pdf of mean voltage, estimated from simulations of 1000 realizations of neurons with ∼ 2000 excitatory and inhibitory pre-synaptic neurons in total, for each neuron. Solid line: fit to a Gaussian distribution. **D**. Top: Voltage heterogeneity against the number of synaptic connections. Bottom: SD of single neuron fluctuations against the number of synaptic connections. Gray: all synapses are on the soma. Black: synapses are distributed on the dendritic tree. Synaptic conductance in C-D was 1nS. **E**. Modeling a network of point neurons that incorporate the location of the synapse along the dendritic tree (see main text, Methods and Fig.S6 for details). (i) Measuring the current and conductance changes at the soma of a simulated layer 5 pyramidal neuron, when activating a synapse on a dendrite. For each synapse we extract the attenuation (*α*) and visibility (Γ) parameters. (ii) An extended-like point neuron. The thicker the synapse is the larger its *α*, while the visibility parameter is color coded. (iii) The empirical distribution of {*α*, Γ}, estimated for each cell type, is then used to simulate a network model of spiking neurons that interact through Eq.(12). **F**. Top: Attenuation vs. the distance of the synapse from soma for the cell in (E). Red line: fit to an exponential decay. Bottom: Same as top, but for the visibility parameter. Note that the decay rate of the visibility is around twice the decay rate of the attenuation parameter.

Encouraged by the single neuron simulations, we introduced a phenomenological *network* model of *‘extended-like’* point neurons that incorporated the effect of distributing synapses on the dendrites (Methods). It is known that changes in the neuronal conductance and currents, as measured at the soma, depend on the proximity of the synapse to the soma ([Rall, 1964, Jack et al., 1975, Koch et al., 1990, Destexhe et al., 2003], Fig.S7). We thus assumed that pre-synaptic action potentials affected the single neuron dynamics in two ways. The first was by changing the current arriving to the soma:

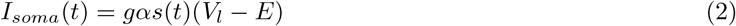

where, as before, *s*(*t*) was a filtered version of the pre-synaptic spikes due to the synaptic time constant, *V*_*l*_ was the reversal potential of the leak current, *g* the synaptic strength and *α* was the *attenuation parameter* that modeled the decay in current change with the distance to the soma. The second contribution was an effective change in the neuronal conductance:

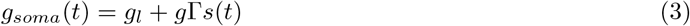

with the leak conductance *g*_*l*_. The *visibility parameter*, Γ, captured the change in somatic conductance as a result of opening of the synapse on the dendrite [Koch et al., 1990]. Thus, in this extended-like neuronal model all synapses were placed on a somatic compartment, but it effectively accounted for their location on the dendrites by changing their visibility and attenuation parameters. The case in which Γ = 1 corresponds to a pure ‘conductance-based model’ of a synapse, while if Γ = 0 the activation of a synapse induces a change in somatic currents but not in the neuron input conductance (‘current-based model’) [Vogels and Abbott, 2005, Burkitt, 2006].

We thus expect that as Γ decreases the clamp of the voltage membrane potential to the effective voltage becomes looser. As a consequence, the time-average membrane potential becomes more heterogeneous across neurons. To show this, we investigated networks of spiking extended-like neurons and analyzed the effect of changing the visibility parameter (Fig.S3). For simplicity, we assumed the same values of the visibility and attenuation parameters for all the neurons. Simulating networks with different levels of Γ, while maintaining the average firing rates of the excitatory and inhibitory populations, we found that for smaller visibility parameter the mean-voltage distribution was wider (Fig.S3B-C).

We next estimated the parameters *α*, Γ for each synapse by directly measuring the change in somatic conductance and currents upon activating a synapse on a dendrite in a multi-compartment neuronal model, and comparing it to the same variations in the phenomenological point model (Fig.4E-F, Fig.S6-7). This procedure is explained in detail in Methods and is illustrated in Fig.4E and Fig.S6A. We found that the average value of the visibility parameter, as estimated from the layer 5 pyramidal neuronal model, was in the range of values that could account for the large *CV*_*ISI*_ and heterogeneous mean voltages we observed in ALM (Fig.S3D-F; Fig.S4).

Further investigation of the distributions of *α*, Γ showed that they depended on the morphology of the cells. For example, the estimated values of *α* and Γ for inhibitory Basket cells, which are electronically more compact than pyramidal neurons, were larger with respect to the estimated parameters in a simulated pyramidal neuron (Fig.4F and Fig.S6D). Importantly, while indeed the visibility and attenuation parameters decayed with the distance of the synapse from the soma, the visibility was always lower than the attenuation (Fig.S6D-E, Fig.4F, Fig.S7C). As a result, while synapses that were sufficiently far from the soma did not contribute much to the shunting of the soma, they still affected the total somatic current (see Methods) and were thus contributing to sub-threshold voltage heterogeneity. This effect can be understood through the passive cable theory. For example, in an infinite cylinder both the visibility and the attenuation parameters decay exponentially with the distance to the soma, but the former decays twice as fast than the latter (Fig.S7C). Intuitively, this is because a current that is injected at the soma (to measure the conductance change) must propagate to the synapse and then back to the soma [Koch et al., 1990].

### Networks of extended-like neurons that operate in a fluctuation-driven regime can account for the supra- and sub-threshold statistics of ALM neurons

Equipped with the estimated distributions of *α*, Γ for a layer 5 pyramidal neuron and for a basket cell, we next simulated a network model of point neurons that interacted through the effective synapses (Eq.(2)-(3) and Eq.(12)). The attenuation and visibility parameters were randomly selected from the joint distribution, *P* (*α*, Γ), which we estimated using the multi-compartmental models (Fig.4F, Fig.S6D). Similarly to networks that neglected the neuronal morphology (Fig.3), the spiking statistics of the extended-like neurons resembled ALM activity (Fig.5B-C). However, in these models, in contrast, the sub-threshold voltage distribution was wider (Fig.5D and compare Fig.5I and Fig.3E) and resembled the sub-threshold voltage statistics of ALM neurons (Fig.1E).

**Figure 5:**
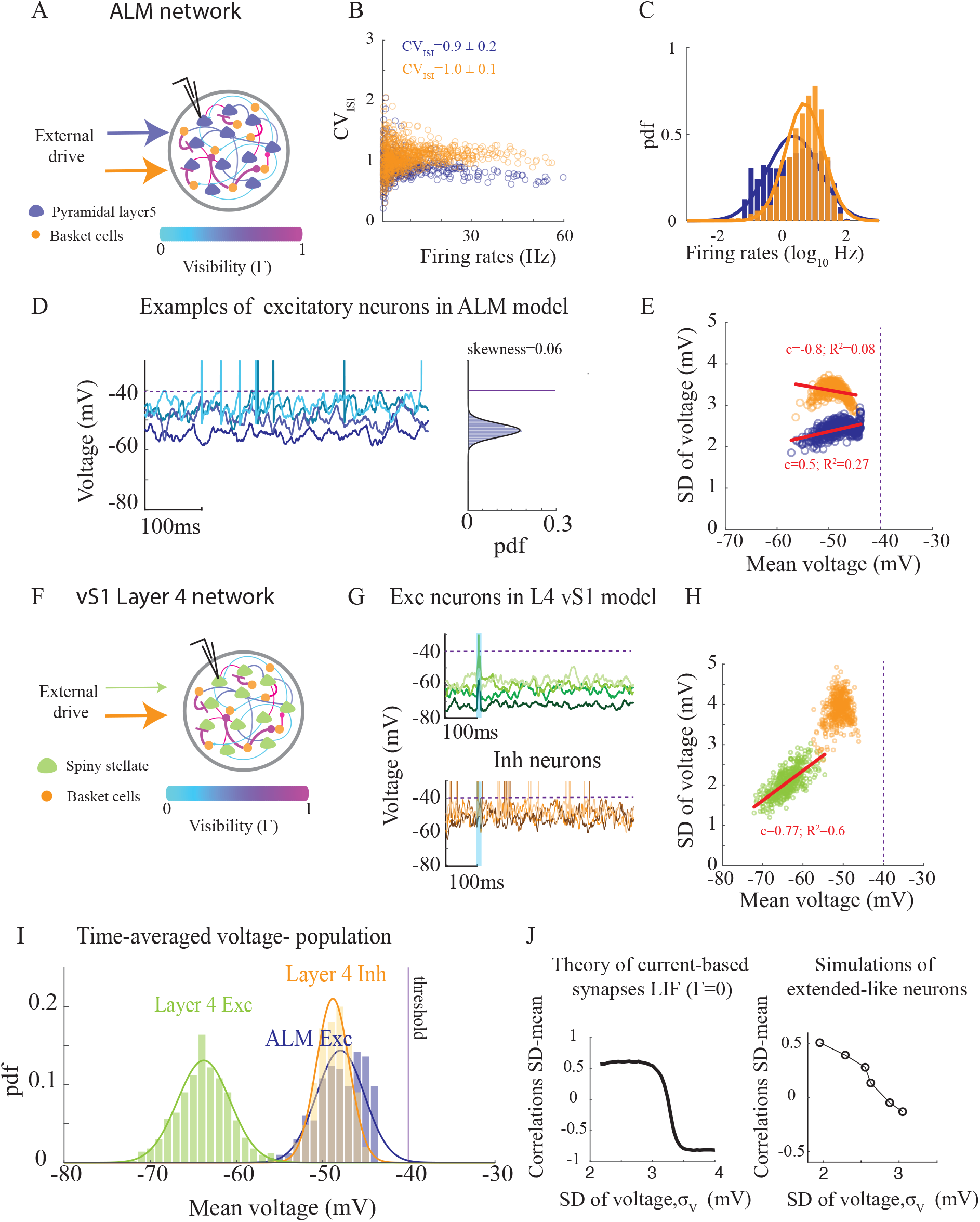
Supra- and sub-threshold statistics of neurons in networks of extended-like point neurons are consistent with the data. **A**. Diagram of an ALM network model with attenuation and visibility parameters estimated from multi-compartment models of layer 5 pyramidal (blue) and Basket (orange) cells. **B**. *CV*_*ISI*_ vs. firing rate in the network for excitatory (blue) and inhibitory (orange) neurons. **C**. Distribution of the log-rates in the network is well-approximated by a Gaussian distribution (solid lines). **D**. Left: Activity of four example neurons. Right: Sub-threshold voltage distribution (excluding spikes) of one of the neurons. Solid line: fit to a Gaussian distribution. Purple line: neuronal threshold. **E**. Voltage SD against mean voltage across the population for excitatory (blue) and inhibitory (orange) neurons. Red: linear fit with a slope (c; mean-SD correlation) and goodness-of-fit (*R*^2^). **F-H**. Same as (A) and (D-E), but for a layer 4 vS1 network. The difference with the network in (A) is in the weak external drive to the excitatory population, which lead to an unbalanced excitatory population. Excitatory neurons fired as a result of short 10*ms* synchronous external drive to all neurons (cyan in G). Note that the excitatory neurons in the model fire only as a result of the synchronous drive. **I**. Probability density function (pdf) of time-average voltage for the neurons in the networks. **J**. Left: Theoretical predication for the correlation between SD and mean voltage across the population in the case of current-based (Eq. (12), Γ = 0) integrate-and-fire neurons, plotted against the voltage SD (Eq.(21)-(22)). Right: Same as left, but for simulations of extended-like networks, as in (A-E). Each point corresponds to a network simulation with network parameters that keep the rate constant, but varies the voltage SD of the neurons.

Specifically, neurons were no longer clamped to an effective reversal and a distribution of time-average voltage in the network was observed. Furthermore, similarly to pyramidal neurons in ALM, the mean voltage and SDs across the excitatory population in the network were positively correlated (compare blue circles in Fig.5E with Fig.1G). This was in contrast to the inhibitory neurons that exhibited a higher level of voltage fluctuations, which were slightly negatively correlated with their mean voltage. In fact, their activity was similar to the fast-spiking putative inhibitory neurons in vS1 (Fig.2G).

This difference in correlations of mean voltage and SDs for excitatory and inhibitory neurons can be understood through a threshold effect. In particular, we derived the sub-threshold statistics for the limiting case of purely current-based models (Γ = 0) and found that these correlations originated from a non-linear effect that depended on the level of fluctuations in the inputs to the neurons (see Methods). The correlations were positive in a low current noise regime, and negative for larger noise levels (Fig.5J). Interestingly, due to the strong recurrent inhibition in the network, the current fluctuations to an inhibitory neuron in our simulations were larger than the fluctuations to an excitatory neuron. As a result, we observed a positive correlation for the excitatory population, but a negative or small correlation for the inhibitory population (Fig.5E).

Finally, by introducing a structured recurrent motif to the unstructured recurrent connectivity that supported bistability in the network (see [Lebovich et al., 2019] and [Inagaki et al., 2019], Supplementary materials), the network was also able to generate selective responses to licking direction during a delay period, without relying on external stimuli (Fig.S5D). Due to the combination of randomness and structure in the connectivity, neurons in the network model, as in the data, also exhibited heterogeneous selective responses in both their supra- and sub-threshold responses (Fig.S5E-F).

### Differences in external drive can explain variations in average membrane potentials of excitatory neurons in ALM and of vS1

We showed that a network model consisting of extended-like point neurons could explain the supra and sub-threshold statistics of ALM neurons. Layer 4 excitatory neurons in vS1, on the other hand, were much more hyperpolarized than ALM neurons. We next investigated the reason for these differences in the resting state of neurons in the two populations.

A majority of the recorded excitatory cells in ALM were layer 5 neurons that are mostly pyramidal, while in layer 4 of vS1 these are spiny satellite cells. Therefore, variations in the distance of the neurons from their threshold in vS1 and ALM could potentially result from morphological differences between these two populations. However, although their morphology differs significantly (Fig.4A, Fig.S6C), we found that the estimated average visibility parameter for spiny stellate neurons and pyramidal neurons was not dramatically different (average visibility of 0.26 and 0.23 for spiny stellate and layer 5 pyramidal neurons, respectively). Indeed, the statistics of sub-threshold activity was not very different than the statistics in the ALM network when we simulated the same network of Fig.5A, but with the estimated parameters for spiny stellate cells from Fig.S6C-D instead of layer 5 pyramidal cells (Fig.S8).

Instead, we hypothesized that the two excitatory populations operated in different dynamical regimes that originated from variations in their network connectivity. Specifically, we conjectured that while ALM operated in a fluctuation-driven regime, in which both excitatory and inhibitory sub-networks were balanced, layer 4 of the barrel cortex was only *partially-balanced*. In this regime, the external excitatory currents and the recurrent inhibition into the inhibitory population are balancing each other, but those for the excitatory population are not.

In the case of two populations (with Γ = 0), reducing the external drive to the excitatory population below a certain value leads to a quiescent excitatory population while maintaining the excitatory-inhibitory balance for the inhibitory population (Methods, [van Vreeswijk and Sompolinsky, 1996]). We therefore simulated a network with similar parameters as the ALM network in Fig.5A-C, but reduced the external drive to the excitatory population (Fig.5F, Methods). As a result, excitatory neurons in the model were hyperpolarized and unbalanced due to a weak external drive to the excitatory neurons in the model. Their mean voltage was about 20*mV* below their threshold (Fig.5G,I) and, similarly to the data, their voltage mean and SDs were positively correlated (here, in contrast to ALM network, the positive mean-SD correlations were due to the reversal potential of the inhibitory synapses and not the neuronal threshold). As a result, excitatory neurons in the network did not fire unless they received an additional synchronous external input (Fig.5G). Interestingly, the sub-threshold fluctuations of the simulated neurons were well-approximated by a Gaussian distribution, and were positively skewed as in the data only when a synchronous external drive was added (Fig.S9A-B).

Finally, while the excitatory population were quiescent, the recurrent inhibition and the external feedforward excitation to the inhibitory population in the model were still approximately balanced; inhibitory neurons fired at a few Hertz, with an average voltage of −50*mV* (around 10*mV* below threshold; Fig.5G,I). Thus, with this architecture, the supra- and sub-threshold activity of neurons in the network were consistent with the activity of neurons in layer 4 network of vS1 (compare Fig.5B-C with Fig.2E,H-I). This result suggested that, similar to the pyramidal neurons of ALM, uncorrelated spiking activity within layer 4 of vS1 was sufficient to drive the inhibitory neurons to fire. This was not the case for layer 4 excitatory neurons in vS1, which fired due to strong synchronous drive.

## Discussion

Spiking activity in the cortex is irregular and heterogeneous across neurons. Intrinsic recurrent network dynamics can account for these features by hypothesizing that excitatory and inhibitory inputs to a neuron are strong but they approximately balanced [Shadlen and Newsome, 1998, van Vreeswijk and Sompolinsky, 1996, Vogels et al., 2005]. The net input current and its fluctuations are then of the same order. As a result, in this fluctuation-driven regime, fluctuations in synaptic inputs can drive neurons to fire action potentials irregularly with spike rates that vary across neurons [van Vreeswijk and Sompolinsky, 1996, van Vreeswijk and Sompolinsky, 1998, Brunel, 2000]. Does the ‘balance hypothesis’ also explain the voltage statistics of the neurons?

We investigated this question by focusing on the anterior lateral motor cortex (ALM), as well as on the vibrissal somatosensory area (vS1). Combining modeling and analysis of intracellular recordings in mice performing decision-making tasks, we found that in ALM the balance hypothesis can explain not only the spiking statistics but also the voltage statistics, provided that one takes into account the morphology of cortical neurons. In contrast, our work suggests that in layer 4 of vS1 the inputs to the excitatory neurons are not balanced, while the fast-spiking inhibitory neurons are.

### Comparison between ALM, vS1 and fluctuation-driven networks

Consistent with previous studies [Softky and Koch, 1993, Hromádka et al., 2008, O’Connor et al., 2010], and in line with previous modeling works [van Vreeswijk and Sompolinsky, 1996, van Vreeswijk and Sompolinsky, 1998, Roxin et al., 2011], neurons in all our data sets exhibited Poisson-like, temporally irregular spike patterns, with a rate distribution that was well-approximated by a log-normal distribution. As expected, network models in which neurons received a relatively large number of pre-synaptic excitatory and inhibitory inputs that were located on the neuronal somatic compartment and balanced each other, could account for the super-threshold spiking statistics of the recorded cortical neurons. These models failed, however, in explaining the large sub-threshold voltage heterogeneity. This was because excitatory and inhibitory inputs induced a large increase in the neuron input conductance; this led the voltage of neurons in the model to fluctuate around an effective reversal potential close to threshold. The value of these reversal potentials was nearly equal for all the neurons. This was in contrast to our analysis of intracellular recordings. Alternatively, in real neurons, synapses are distributed along the dendritic tree. As we showed, synapses that are farther from the soma lead to current flow into the soma without a large increase in the neuronal conductance (Fig.4F, Fig.S7C). Therefore, incorporating an effective morphology of cells into the network allowed to capture the heterogeneity in the means and standard deviations of the membrane potentials across neurons in the network, as well as the correlations between them (Fig.5E,H,J).

Average membrane potentials of ALM excitatory neurons and vS1 fast-spiking (FS) inhibitory neurons were typically around 10 − 12*mV* below their threshold, which together with their fluctuations and their rate distribution was consistent with the hypothesis that the excitatory and inhibitory inputs to these populations were approximately balanced. Similar to the ALM network, the mean voltages of excitatory neurons in vS1 was also heterogeneous, and their sub-threshold fluctuations were comparable to those in ALM neurons (average SD of 2 − 3*mV*). However, excitatory neurons in vS1 were much more hyperpolarized compared to ALM neurons.

In fact, excitatory neurons in layer 4 vS1 network (L4E) were so far from their threshold that they barely spiked. This suggested that their excitatory inputs were not strong enough to balance their inhibitory inputs. On the other hand, we found that the distribution of mean voltages of the FS neurons in vS1 was very similar to the one of excitatory neurons in ALM. We thus posit that in the inhibitory population in layer 4 vS1 network the feedforward excitation was approximately balanced by the mutual inhibition.

We have shown in our simulations that a spiking neuronal network can operate in such a *partially-balanced* regime if the ratio between the feedforward drive to the excitatory and inhibitory populations was too small with respect to the recurrent interactions ([van Vreeswijk and Sompolinsky, 1996, van Vreeswijk and Sompolinsky, 1998] and Methods). We thus posit that the thalamic excitation to inhibitory neurons was significantly stronger than for excitatory neurons. This hypothesis is consistent with experimental results on the thalamic input to layer 4 neurons in vS1 cortex [Gabernet et al., 2005, Cruikshank et al., 2007, Gutnisky et al., 2017].

Finally, in contrast to L4E neurons, layer 5 excitatory neurons in vS1 (L5E) spiked at similar rates as ALM neurons (a majority of which were also layer 5 neurons). Consistent with this observation, the mean voltages of these layer 5 excitatory neurons were closer to their thresholds than L4E neurons. However, they were on average still more hyperpolarized than ALM neurons. With respect to the model, we note that a clear distinction between a fluctuation- and a non-fluctuation-driven regime (or a fully- and partially-balanced regime) is well-defined only in the theoretical limit of an infinite number of inputs per neuron. In finite networks, in which the balance can only be approximated, the transition between these two regimes becomes less clear. In this sense, L5E neurons might operate on the border of the two regimes such that recurrent inhibition is only partially-balanced by the feedforward excitation.

### Relations to previous studies

Consistent with previous recordings in auditory [DeWeese and Zador, 2006] and visual [Tan et al., 2014] cortices, we found that the membrane potentials of neurons in vS1 were more positively skewed than those of excitatory neurons in the ALM and inhibitory FS neurons in vS1 (Fig.S2). However, the observation of deviations from Gaussianity of the membrane potential of cortical neurons (measured, e.g., by the skewness of the distribution) is not *per se* an unequivocal test of the balance hypothesis. This is because, although it predicts that synaptic currents to a neuron should be approximately Gaussian, the distribution of the voltage can be positively or negatively skewed due to proximity to threshold, to the reversal of chloride channels, or other non-linear effects (Fig.S9).

A stronger test to assess this hypothesis is provided by the distribution of the mean voltage (or distance to threshold) across the population. As previously reported in visual [Tan et al., 2014] and auditory [DeWeese and Zador, 2006] cortices, the voltage of excitatory neurons exhibits synchronous fluctuations, about 20mV below threshold during spontaneous activity. This suggests that, at least in this condition, the neurons are driven by strong and synchronized external synaptic inputs [Tan et al., 2014]. This is similar to our findings in layer 4 excitatory neurons of vS1, which we suggest operated in a partially-balanced regime.

Comparing sub-threshold voltage dynamics in models of recurrent networks with recorded neurons requires extra care in the way synaptic interactions are modeled. Previous studies showed that cortical-like spiking dynamics emerge in networks of neurons interacting through realistic synapses, in which a pre-synaptic spike leads to a transient change in the post-synaptic conductance, [Hertz et al., 2003, Vogels and Abbott, 2005, Sanzeni et al., 2022]. However, in contrast to our network models that account for the proximity of the synapses to the somatic compartment, in these previous studies all synapses influenced the soma in the same way; we showed that this leads to small voltage heterogeneities across neurons in the network. A possible way to increase the voltage heterogeneity in these networks of one-compartment neurons is to consider wider distributions of the in-degree connectivity [Landau et al., 2016, Sanzeni et al., 2022]. However, we found that incorporating such large heterogeneous in-degree distributions in our networks that fit well the activity of ALM neurons, only slightly increased the voltage heterogeneity before the firing rate distribution started to deviate significantly from its log-normal shape.

### Perspectives

An interesting future direction would be to investigate our model, which combines a mixture of conductance and current synapses in the mean-field limit of a large number of connections. Indeed, one of the problems in a model of conductance-based synapses that are all localized on the soma is that by using the synaptic scaling of balanced networks to keep the current fluctuations finite, it makes the neurons very leaky due to the very large conductances. Alternatively, in a model with a mixture of current and conductance synapses that mimics the distribution of synapses on the dendrites, one can obtain a finite conductance and finite current fluctuations by scaling the average visibility of the synapses, Γ, instead of scaling directly the connection strengths. Specifically, scaling Γ with the inverse of the number of pre-synaptic connections in the network (and thus *α* with the inverse of the square root of the number of pre-synaptic connections) guarantees that both the effective neuronal conductance and current fluctuations are finite for large networks. Analytical study of such scaling is outside the scope of this paper.

We studied whether fluctuations in a cortical network were consistent with a mechanism in which they emerged from the recurrent network dynamics. In reality, membrane fluctuations are a combination of external and recurrent fluctuations, and the relative proportion can vary between areas, populations and behavioral states. Therefore, some of the variance in voltage fluctuations could be potentially attributed to non-local recurrent connections, as we argued, for example, for vS1 neurons. It would be interesting to control for such unobserved external inputs for local cortical networks while measuring the neuronal membrane potential. This will potentially require multi-brain area recordings with voltage measurements to determine the respective contributions of feedforward and recurrent connectivity to membrane potential fluctuations of neurons.

There are other, non-inclusive, mechanisms that can contribute to the increase of voltage heterogeneity that we did not include in our network models. For instance, we modeled the effect of activating simple AMPA/GABA synapses in multi-compartment models on the soma, but cortical neurons consist of non-linear NMDA synapses [Nevian et al., 2007, Larkum et al., 2009], voltage-gated mechanisms that affect the input resistance [Li et al., 2020], as well as active non-linear ion channels that can lead to localized spike-like activity [Larkum et al., 1999, Waters et al., 2003]. Incorporating such active channels in network models is an important future direction.

### Functional implications

Finally, we speculate why one cortical region is more fluctuation-driven than another. We suggest that a partially-balanced operating regime might be beneficial for detection tasks. In this case, excitatory neurons that are hyperpolarized will be insensitive to external and internal fluctuations and this could reduce the false-positive detection rates. They will also be less prone to weak, but synchronous, variations in the external drive if the inhibitory neurons are balanced. Indeed, in this regime the inhibitory neurons react very fast to external stimuli, as there are many neurons that are ready to fire (a phenomenon known as ‘fast tracking’ [van Vreeswijk and Sompolinsky, 1996]). This will keep the excitatory population far from their threshold, unless they receive a synchronous drive that, at least instantaneously, will be stronger than the external drive to the inhibitory neurons. This might be achieved through a ‘window of opportunity’, based on a delay between the two populations [Yu et al., 2016, Gutnisky et al., 2017]. This type of operating regime might be suitable for areas involved in stimulus detection, such as the somatosensory and the auditory cortices.

In contrast, strong and balanced excitation and inhibition may be needed for neurons to generate selective responses that are robust to noise or generate stable memory states [Rubin et al., 2017], as in ALM [Inagaki et al., 2019]. Furthermore, neurons that are driven by fluctuations in their synaptic inputs are also very sensitive to heterogeneities in the network activity and connectivity [Hansel and van Vreeswijk, 2012, Pehlevan and Sompolinsky, 2014, Lebovich et al., 2019, Darshan et al., 2017]. This feature is beneficial in networks that undergo synaptic reorganization, as it allows the neurons to develop task-related activity during learning with minimal synaptic modifications [Kim et al., 2022].

In the near future, advances in technologies will allow simultaneous recording of the subthreshold activity of many cortical neurons (e.g., [Abdelfattah et al., 2019, Adam et al., 2019]). Understanding the statistics of these measurements and how they vary with the dynamical regime of the cortex will be an important step towards deciphering the operating regime of cortex and its functional implications.

## Acknowledgements

We would like to thank Larry Abbott, Sandro Romani and Alessandro Sanzeni for their valuable feedback. This work was supported by the Howard Hughes Medical Institute.

## Methods

### Data analysis

Spiking statistics for ALM neurons were analyzed based on extracellular recordings using silicon probes of 667 putative pyramidal neurons and 74 putative fast-spiking neurons [Inagaki et al., 2019]. Inter-spike-intervals (ISI) were calculated for the delay or sample periods in each of the trials and were combined together in order to calculate the coefficient of variation of the inter-spike-intervals (ISI) of the neurons, which were calculated as the standard deviation of the ISI divided by the mean of the ISI distribution.

Spiking statistics for vS1 neurons were analyzed in a similar way. They were based on juxtacellular recordings of 95 regular spiking neurons and 43 putative fast-spiking neurons in layer 4 [Yu et al., 2016], and 53 regular spiking neurons and 22 putative fast-spiking neurons in layer 5 [Yu et al., 2019], with some of the fast-spiking neurons that were also identified as inhibitory neurons through optogenetic tagging [Yu et al., 2019]. Inter-spike-intervals (ISI) were calculated for each non-whisking epoch and were combined together in order to calculate the coefficient of variation of the inter-spike-intervals (ISI) of the neurons.

Membrane potentials of ALM and vS1 neurons were obtained based on whole-cell recordings. The recording details for ALM neurons are given in [Guo et al., 2017, Inagaki et al., 2019] and for vS1 neurons in [Yu et al., 2016]. For ALM neurons, we combined two datasets from auditory and tactile delayed response task (crosses and circles in Fig.1F-G, respectively). We analyze the membrane potential of neurons that exhibited stationary membrane potentials during the delay period (47/89 recorded neurons). For vS1 we had 37 L4E neurons and 38 L5E neurons. The mean voltage of the five FS interneurons of layer 5 and the eight layer 4 FS interneurons in vS1 were similar and we thus combined them for the analysis.

Spike threshold was defined as the moment when *dV/dt* first crossed 33% of its maximal value during the trial [Yu et al., 2016]. Spikes were removed by interpolating the pre-spike (-15 ms for ALM neurons, 0.5ms for vS1 neurons) and post-spike (15 ms and 10ms, respectively) voltage.

Mean and standard deviation (SD) of threshold were then calculated for each cell that spiked and mean and SD of the population threshold was calculated based on the cell-specific mean thresholds. To obtain the distance of ALM neurons to their threshold, we subtracted the mean threshold from the mean voltage of each neuron. For neurons that did not spike, we used the minimal threshold across all recorded ALM neurons as their spike threshold.

We then calculated the moments of the single-cell voltage distribution. Specifically, we concatenated all the non-whisking periods for neurons in the vS1, or during delay response periods for ALM neurons, after we subtracted the mean voltage at each epoch/period. We then calculated the SD and the skewness of these concatenated and mean-subtracted voltage traces for each neuron. To calculate the mean activity, we averaged the mean values of all epochs/periods. For neurons in vS1 we also corrected for possible drift in the mean voltage by first regressing the mean voltage per trial against the trial number, and then subtracting the slope of the regression from the voltage trace.

### Limitations in estimating voltage heterogeneity in the data

We found that the time-average voltage of neurons had a wide distribution across neurons, with SD that could go up to 6 − 8mV. We note that this estimate was probably an upper bound for the real voltage distribution for several reasons; 1) Although we tried to control for the state of the network, for example by analyzing only non-whisking epochs or delay periods, the network state could still change over the course of recordings and between subjects. 2) There were inherent biases in the recordings. For example, in some cases the mean voltage was drifting over time. While we corrected for the drift, this might have been imperfect. 3) We used blind recordings, which were presumably targeting the soma, however it might be that in some cases the recordings were at dendrites or axons.

### Mapping synapses on dendrites to synapses on an extended-like point neuron

#### Multi-compartment single neuron model

We used NEURON [Carnevale and Hines, 2006] (Fig.4, Fig.S6, Fig.S7) to simulate multi-compartmental models from the morphological reconstructions of three different cells from the mouse. motor cortex layer 5 pyramidal cell, visual cortex layer 4 basket cell, and somatosensory layer 4 spiny stellate cell [Scala et al., 2021, Scala et al., 2019, MacLean et al., 2005] (neuromorpho IDs: NMO161366, NMO130658 and NMO02484 respectively).

Input resistances differ between *in vitro* to *in vivo* conditions, presumably due to the constant synaptic load which is much lower in a slice. For estimating *α*, Γ we simulated *in vivo* conditions, assuming that the effect of a single synapse should be measured while others are active. In contrast, for exploring the voltage distribution in Fig.4, we simulated *in vitro* conditions, where no synapses were active. We then activated the synapses by simulating pre-synaptic Poisson neurons with firing rates sampled from the log-normal rate distribution of the pyramidal and fast-spiking ALM neurons.

For in vitro conditions the specific membrane resistivity (Rm) for the pyramidal cell was set to 8, 000Ω*cm*^2^, for the basket cell to 14, 000Ω*cm*^2^, and for the spiny stellate cell to 10, 000Ω*cm*^2^. This yielded an input resistance (Rin) of 164, 141, 302*M* Ω, respectively. These values are within the experimental range measured *in vitro* [Oswald et al., 2013, Lefort et al., 2009, Dougherty et al., 2014]. For *in vivo* conditions, Rm for the pyramidal cell was set to 3500Ω*cm*^2^, for the basket cell to 3000Ω*cm*^2^, and for the spiny stellate cell to 1000Ω*cm*^2^. This yielded an input resistance (Rin) of 57, 36, 45*M* Ω, respectively. These values are within the experimental range measured *in vivo* [Bindman et al., 1988, Paré et al., 1998, Guo et al., 2017]

All multi-compartmental models had an axial resistance (Ra) of 100 Ω*cm*, and specific membrane capacitance (C) of 1*μF/cm*^2^. Unless specified otherwise, synaptic conductance in the models was set to 0.5nS, with reversal potential of 0mV and −80mV for the excitatory and inhibitory synapses, respectively.

#### Estimating the change in somatic conductance and somatic current upon activation of one synapse in a multi-compartment model

Opening of a synapse on a dendrite results in variations in the neuronal conductance, as measured at the soma, as well as the current flowing into the soma. The effectiveness of these changes reduces as a function of the distance of the synapse from the soma. Our goal was to construct a one-compartment model of the soma, in which all synapses are located on the same compartment, but that also effectively accounts for their location on the dendrites. To this end, we used a multi-compartment model and opened a synapse on a dendrite while measuring the conductance and current changes at the soma. We then constructed a one-compartment model and parameterized its synapses such that activating a synapse on the the neuronal compartment would result in the same amount of conductance and current changes as the synapse in the multi-compartment model

Specifically, let us consider a synapse of conductance *g* located on the dendrite of a neuron. We first measured the somatic voltage change, 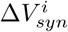, upon activating the *i*’th synapse on the dendrite (static response; Fig.S6A). We then measured the somatic voltage change, 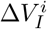, in response to an additional current step injected at the soma, *I*. We calculated the somatic conductance:

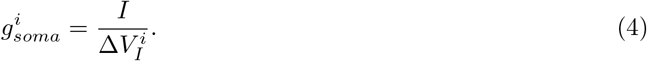

and defined the visibility of the synapse that captured the change in somatic conductance, normalized by the maximal possible change [Koch et al., 1990]:

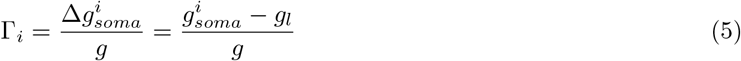

with the neuronal conductance, *g*_*l*_, measured before the activation of the synapse.

We next approximated the current arriving at the soma upon activating the synapse on the dendrite, 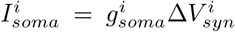, and defined the attenuation parameter, *α*_*i*_, which approximated the decay in current, normalized by the maximal possible current that the *i*’th synapse could inject as:

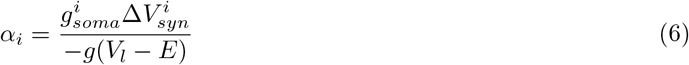

Here, *E* is the reversal potential of the synapse and *V*_*l*_ the reversal potential of the leak current. Both the visibility and attenuation parameters decay with the electrotonic distance of the synapse from the soma (Fig.S7A-C).

We determined the spatial dependence of the attenuation and of the visibility parameters by performing this calculation for each synapse while changing the location of the synapse on the dendritic tree. This provided us with the empirical distribution of {*α*_*i*_, Γ_*i*_} for each of the multi-compartment model considered. Importantly, we found that the visibility parameter decayed with the distance of the synapse from the soma much faster than the attenuation parameter. For example, the visibility parameter in an infinite cylinder with small synaptic conductance followed Γ(*x*) ∝ *e*^*−*2*x/λ*^, with *λ* the space constant and *x* the distance of the synapse from the soma. On the other hand, the attenuation parameter decayed more slowly, *α*(*x*) ∝ *e*^*−x/λ*^ (Fig.S7C, [Koch et al., 1990]).

#### An extended-like point neuron model

For the effective one-compartment model, we considered a neuron in which its membrane potential followed:

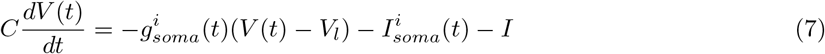

with *C, V*_*l*_ being the capacitance and reversal membrane potential, *I* was the current injected into the compartment, and where we omitted the post-synaptic index to simplify notations.

We modeled the change in neuronal conductance due to opening of the *i*’th synapse to be linear in the synaptic conductance:

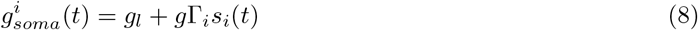

with the leak conductance, *g*_*l*_. The change in synaptic conductance induced by a pre-synaptic spike was *g*, and *s*_*i*_(*t*) was a filtered version of the pre-synaptic spike. The visibility parameter, Γ_*i*_, was estimated from the multi-compartment model (see above).

The effective change in current arriving at the compartment due to the opening of a synapse was modeled as:

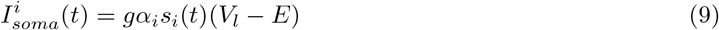

with *E* being the synaptic reversal potential. The attenuation parameter, *α*_*i*_, which modeled the decay in somatic potentials when activating synapses that are far from the soma, was estimated from the multi-compartment model.

This choice of modeling guaranteed that opening the *i*’th synapse with its estimated *α*_*i*_, Γ_*i*_ in the effective one-compartment model and on the dendrite of the multi-compartment model would lead to comparable changes (at least at steady state) in somatic currents and conductance in both models (Fig.S7B,D,F).

Inserting Eqs.(8) and (9) into Eq.(7), rearranging, and omitting the synaptic index to simplify notations yields:

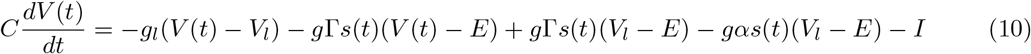

We next wrote Γ = *αρ* and rewrote Eq.(10) as:

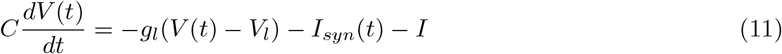

where

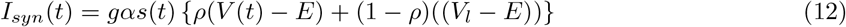

With this formulation, the parameter *ρ* interpolated between a fully ‘conductance-based’ model of the synapse (*ρ* = 1), in which a pre-synaptic spike led to a transient change in the post-synaptic conductance, and a fully ‘current-based’ model (*ρ* = 0), in which post-synaptic conductance was independent of pre-synaptic spikes [Vogels and Abbott, 2005, Burkitt, 2006, Hansel and van Vreeswijk, 2012]. We thus refer to *ρ* as the *mixing parameter*.

Finally, we note that the average mixing parameter that we estimated using the multi-compartment model only weakly depended on *g*. However, the average attenuation parameter decreased with *g*. Yet, any change in the average attenuation could be compensated through increasing the strengths of the synapses in our network models in a way that best fit the sub-threshold fluctuations. It is thus sufficient to estimate *α, ρ* from the multi-compartment model only for one fixed value of *g*.

#### Spiking neuronal network

We consider a network of *N*_*E*_ excitatory and *N*_*I*_ inhibitory neurons randomly connected. We denote by Λ the adjacency matrix of the network connectivity, defined as 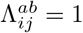 with probability *p*_*a*_ = *K*_*a*_*/N*_*a*_ and 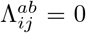 otherwise. Here, *a, b* ∈ {*E, I*} and *i, j* = 1…*N*_*a*_. Thus, each neuron thus receives, on average, *K*_*E*_ and *K*_*I*_ synaptic inputs.

Neurons are modeled as leaky integrate-and-fire elements. The membrane potential of neuron 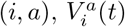, obeys

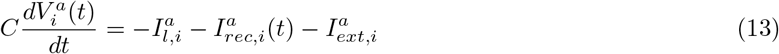

where *C* is the capacitance, 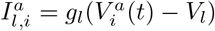 is the leak current whose reversal potential is *V*_*l*_ and *g*_*l*_ the leak conductance. Whenever the voltage reaches a threshold, 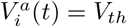, it is reset to 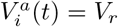.

Following the reduction procedure explained above, the recurrent input, 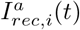, into neuron (*i, a*) is modeled as

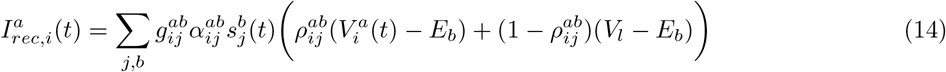

with *E*_*b*_ being the synaptic reversal potential and 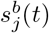:

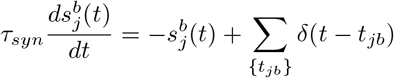

Here, *τ*_*syn*_ is the synaptic time constant and the sum is over all spikes emitted by neuron (*j, b*) at times *t*_*jb*_ *< t*.

As in Eq.(11), the parameter 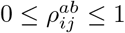 interpolates the synapses from being fully conductance synapses 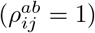 to fully current synapses 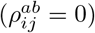. The parameter 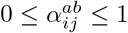 models the attenuation of the synaptic activity as a function of its distance from the soma, with small 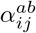 for synapses that are far from the soma. Note that results of multi-compartment simulations shows that 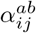 and 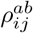 are correlated. Yet, in the homogeneous case, for which 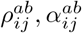 and synaptic strengths are the same for all neurons within a population (Fig.3), we absorb the parameter *α* into the synaptic strengths, *g*_*ab*_.

Finally, we model the external input, 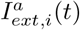, as

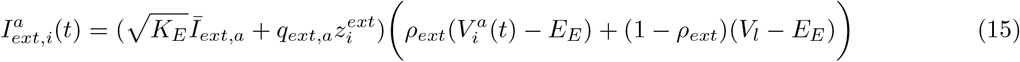

with 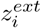 being a Gaussian random variable with zero mean and unity variance and *Ī*_*ext,a*_ is of 𝒪 (1) [van Vreeswijk and Sompolinsky, 1998]. Here, to simplify simulations, we modeled the feed-forward synapses with an average parameter *ρ*_*ext*_ and absorbed the attenuation parameter into the input strength.

#### Choice of network parameters

We chose the parameters of the network connectivity and external drive such that the distributions of neuronal firing rates in the populations fit well the data. For layer 4 model of vS1 we chose parameters such that the excitatory neurons are quiescent, while the inhibitory neurons are not. In a network of current-based synapses (Γ = 0) this is achieved by breaking the inequalities that determine the existence and stability of the balance state. Specifically, in a current-based network a partially-balanced regime, in which the inhibitory neurons are balanced and the excitatory neurons are unbalanced and quiescent, appears when 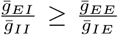 and 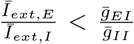 [van Vreeswijk and Sompolinsky, 1998]. The conditions for quiescent excitatory neurons in networks consisting of a mixture of current and conductance-based synapses are more involve. However, we find in simulations that excitatory neurons become silent when decreasing the fraction 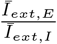. This is the main difference between the parameters of the ALM model network and vS1 layer 4 model network (see Table 1). Interestingly, in the context of the supra-stabilized networks (SSN) [Rubin et al., 2015], this partially-balanced regime is related to the supersaturation phenomenon in which the external drive to the excitation is too small, resulting in a quiescent excitatory population, with active inhibitory neurons [Ahmadian et al., 2013].

**Table 1:** Network simulation parameters for figures in main text.

| Neuron parameters |  | Fig.3 | Fig.5-ALM | Fig.5-S1 |
| --- | --- | --- | --- | --- |
| $dt$ | simulation time step | 0.5 ms | 0.5 ms | 0.1 ms |
| $g_l$ | neuronal conductance | $0.1mS/cm^2$ | | |
| $C$ | neuronal capacitance | $1\mu F/cm^2$ | | |
| $V_{th}$ | spike threshold | -40mV | | |
| $V_r$ | voltage reset after spike | -52mV | | |
| $V_l$ | voltage reset after spike | -55mV | | |
| $E_E$ | reversal potential of exc synapses | 0mV | | |
| $E_I$ | reversal potential of inh synapses | -80mV | | |
| Network parameters |  |  |  |  |
| $\tau_{syn}$ | synaptic time constant | 3 ms | | |
| $N$ | number of neurons | 10000 | | |
| $N_E$ | number of excitatory neurons | $N/2$ | | |
| $N_I$ | number of inhibitory neurons | $N/2$ | | |
| $K_E$ | average number of exc synapses to a neuron | 400 | | |
| $K_I$ | average number of inh synapses to a neuron | 400 | | |
| $p_a$ | connection probability | $K_a/N_a$ | | |
| $\bar{g}_{EE}$ | exc to exc synaptic weight | $0.00133mS/cm^2$ | 0.0026 | 0.0026 |
| $\bar{g}_{IE}$ | exc to inh synaptic weight | $0.02mS/cm^2$ | 0.039 | 0.039 |
| $\bar{g}_{II}$ | inh to inh synaptic weight | $0.133mS/cm^2$ | 0.22 | 0.28 |
| $\bar{g}_{EI}$ | inh to exc synaptic weight | $0.08mS/cm^2$ | 0.156 | 0.225 |
| $\alpha_{ij}^{ab}$ | attenuation parameter | 1 | estimated | estimated |
| $\rho_{ij}^{ab}$ | mixing parameter | 1 | estimated | estimated |
| $\bar{I}_{ext,E}$ | external input to exc | $0.00533\mu A$ | 0.0028 | 0.001 |
| $\bar{I}_{ext,I}$ | external input to inh | $0.00747\mu A$ | 0.006 | 0.006 |
| $q_{ext,E}$ | disorder in external input to exc | $0.008\mu A$ | 0.008 | 0.008 |
| $q_{ext,I}$ | disorder in external input to inh | $0.005\mu A$ | 0.005 | 0.005 |
| $\rho_{ext}$ | average mixing parameter of ext inputs | 1 | 0 | 0 |

To model the synchronous external drive of layer 4 vS1 network we added to 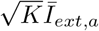. in Eq.(15) another brief external drive of 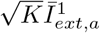, with 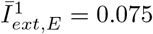 and 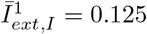, for 10ms.

### Mean-field analysis of sub-threshold statistics of integrate-and-fire neurons

#### Current-based integrate-and-fire neurons

In this section we obtain the statistics of the time-averaged voltage and its SD across neurons in the network. We consider the homogeneous case of a current-based integrate-and-fire neuron (Γ = *ρ* = 0, and we absorb *α* into *g*_*ab*_). We assume that the network is asynchronous and follow standard mean-field approach to describe the total input to a neuron, 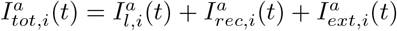, by a Gaussian process:

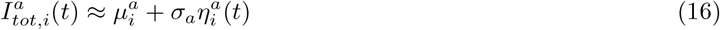

where 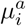 is the time-average input, *σ*_*a*_ is the SD and 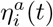 is a white-noise term (assuming *τ*_*syn*_ ≪ *C/g*_*l*_, [Brunel, 2000]). The input SD, *σ*_*a*_, is the same for all neurons in the population (see below). In contrast, the mean input, 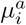, is different across neurons due to variations in the pre-synaptic spike rates and the number of synapses per neuron. In large networks the statistics of this time-average input, which determines the heterogeneity of the mean input currents across the population, is Gaussian. We thus write 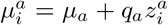, with random variables, 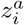, that are drawn independently from a Gaussian distribution with mean 0 and variance 1.

The total input to a neuron of population *a* under this mean-field analysis is thus fully described by three sufficient statistics, *s*_*a*_ = {*μ*_*a*_, *q*_*a*_, *σ*_*a*_}. With this input statistics, the voltage dynamics of a neuron with the inputs of Eq.(16) is reflecting a realization of a typical neuron in the network. Inserting Eq.(16) into Eq.(13) leads to a Langevin equation. The voltage dynamics is thus given by a Fokker-Plank (FP) equation, from which it is possible to calculate the transfer function that relates the mean input and its fluctuations to the mean firing rate [Brunel, 2000, Roxin et al., 2011]:

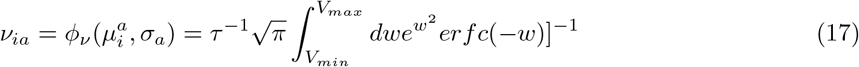

with 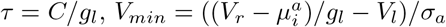 and 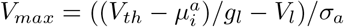.

Using the equilibrium distribution of the FP equation, one can also calculate the moments of the voltage distribution [Hansel and Van Vreeswijk, 2002]. For example, the time-average voltage of a neuron (*i, a*) is:

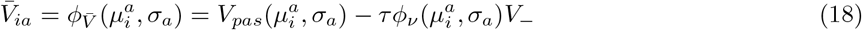

with *V*_*−*_ = *V*_*th*_ − *V*_*r*_ and 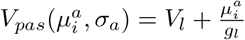. Similarly, the voltage SD yields:

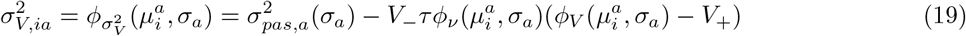

with 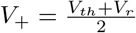 and 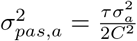.

The equations for the mean and SD of the voltage are simple to understand. The first term in Eq.(18) and Eq.(19) results from bombarding a passive neuron with Poissonian inputs. The second term in these equations results from the spiking mechanism and mainly affects neurons that are close to threshold. It is proportional to the rate of the events (*τν*) and the size of the reset, *V*_*−*_.

Using Eqs.(17)-(19) we can then calculate the mean and the variance of the voltage and rate distributions across the population by averaging over the neurons:

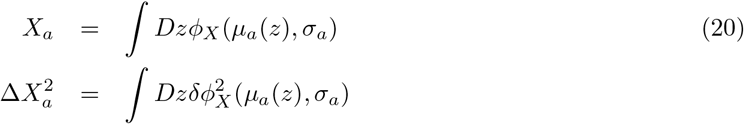

with 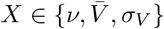 and 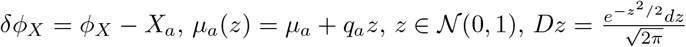.

#### Correlations between voltage mean and SD across neurons in the network

Both in ALM and vS1 data, as well as in our network simulations, we observed that the voltage mean and SDs of neurons can be positively or negatively correlated, depending on the network state. We use the mean field analysis for Γ = 0 to explain this correlation structure.

Equation (19) shows that for neurons that are far from threshold, the voltage SD is governed by the term 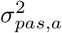, which only depends on the input fluctuations, *σ*_*a*_. Thus, fluctuations of these neurons are the same, independent of their mean voltage. When the mean input current is sufficiently large, both the mean voltage and the SD saturate (due to the threshold). Thus, we expect that the input noise (*σ*_*a*_) will determine the correlations strength (and its sign) between the voltage SD and its mean.

To show the dependence of the mean-SD correlation on the input SD (or similarly, the voltage SD), we fix the mean rates and SDs of the neurons to be in the range of the pyramidal neurons in ALM (6 ± 8*Hz*), and numerically invert Eqs.(20) with *X* = *ν*, while changing *σ*_*a*_ to obtain *μ*_*a*_ and *q*_*a*_. We then used the sufficient statistics, *s*_*a*_ = {*μ*_*a*_, *q*_*a*_, *σ*_*a*_}, to plot the correlations between the mean and voltage SD:

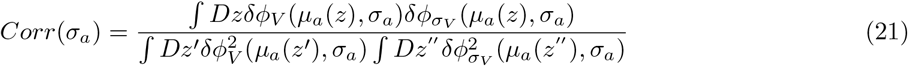

against the average voltage SD (Fig.5J, left), given by

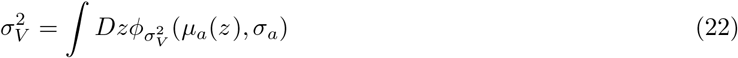

This analysis shows that correlations in the mean-SD voltage across the population is determined by the level of the input SD.

Finally, the inhibitory neurons in our network simulations operate in a regime in which they fire at higher rates than the excitatory neurons. This, together with the strong recurrent interactions to the inhibitory neurons, lead to higher input SD for the inhibitory neurons than for the excitatory neurons in the network (see parameters). As a result, the mean and voltage SDs of the excitatory neurons are positively correlated, while they are less, and even negatively, correlated for the inhibitory population (Fig.5E,J).

#### Conductance-based integrate-and-fire neurons

Here we obtain the statistics of the time-averaged voltage for the homogeneous case of a conductance-based integrate-and-fire neuron (Γ = *ρ* = 1, and we absorb *α* into *g*_*ab*_). In contrast to the case of current-based neurons, for which we only need (*μ*_*a*_.*q*_*a*_, *σ*_*a*_) to describe the distributions of the input to the neurons and thus the rate and voltage distributions of the population, in the case of conductance-based synapses we need six sufficient statistics, *s*_*a*_ = {*μ*_*aE*_, *q*_*aE*_, *σ*_*aE*_, *μ*_*aI*_, *q*_*aI*_, *σ*_*aI*_}. These are the statistics that define the Gaussian input of the total excitatory and total inhibitory synaptic conductances to a neuron (*i, a*):

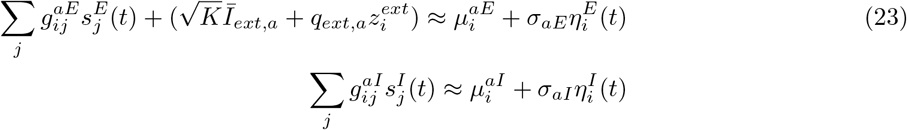

with 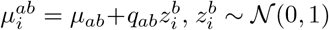 and 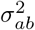 is the variance of the white noise term 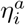, with 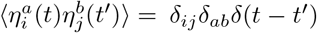. For simplicity of notations we write the statistics in vector notations, *s*_*a*_ = {***μ***_*a*_, ***q***_*a*_, ***σ***_*a*_} and 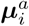.

Similarly to the case of current based neurons, it is possible to use a FP formalism to analytically describe the firing rate and membrane potential distribution of a conductance-based neuron [Richardson, 2004]. We can then use the equilibrium FP distribution to calculate the moments of the distribution (see Supplementary materials). For example, the time-average voltage of a neuron (*i, a*) yields:

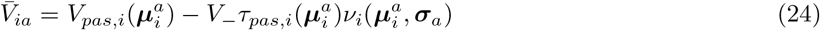

with

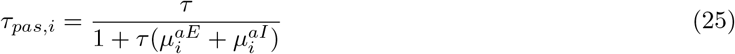

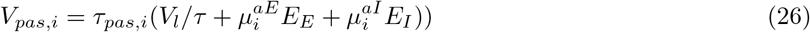

Thus, the mean of the voltage for conductance-based integrate-and-fire neurons have the same form as the mean voltage of the current-based integrate-and-fire neurons (Eq.(24)). It consists of the mean of a passive neuron, bombarded by Poissonian inputs, together with another term that results from the threshold.

Equations (24)-(26) show why the amount of voltage heterogeneity in conductance-based networks are low for large networks. In contrast to networks of current-based integrate-and-fire neurons, for which the currents can be dynamically balanced such that *μ*_*a*_ and *q*_*a*_ are both 𝒪 (1) [van Vreeswijk and Sompolinsky, 1998], for conductance-based neurons there is no balancing of the conductances [Sanzeni et al., 2022]. As a result, the average conductances, ***μ***_*a*_, dominate the quenched disorder, ***q***_*a*_, which due to the central limit theorem is always a 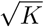 order of magnitude smaller than ***μ***_*a*_. In the large *K* limit, we obtain that to the leading order:

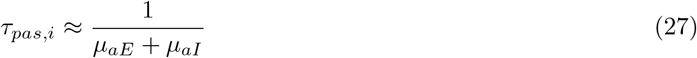

and

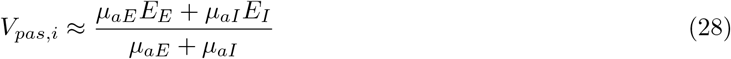

Thus, the first term of Eq.(24) only depends on the mean conductances, ***μ***_*a*_, and the synaptic reversals; it is the same for all neurons in the network. Importantly, this result is independent of the scaling of the connectivity. Heterogeneity in firing rates can, in principle, contribute to voltage heterogeneity due to the second term in Eq.(24). However, it only affects neurons that are close to the threshold, and does not seem to increase the voltage heterogeneity in simulations (Fig.3G). To verify the applicability of Eq.(24) to our network simulations even though *τ*_*syn*_ was not much smaller than *τ* (see Table1), we used the estimated conductances and firing rates from the simulations, together with Eqs.(24)-(26) to calculate the SD of the time-averaged voltage distribution across the neurons (Δ_*V*_). These are the theoretical lines in Fig.3G. This analysis explains why neurons in conductance-based networks are all clamped to more or less the same mean voltage, with a narrow distribution around this number. Similar arguments also hold for the voltage SD.

#### Quantification and statistical analysis

Simulations were done using *C*^++^ and NEURON 7.8. Analyses of simulations and data were done using Matlab (Mathworks), Python and NumPy. Data are presented as mean ± standard deviation (SD), unless otherwise noted.

## Supplementary Figures

**Figure S 1:**
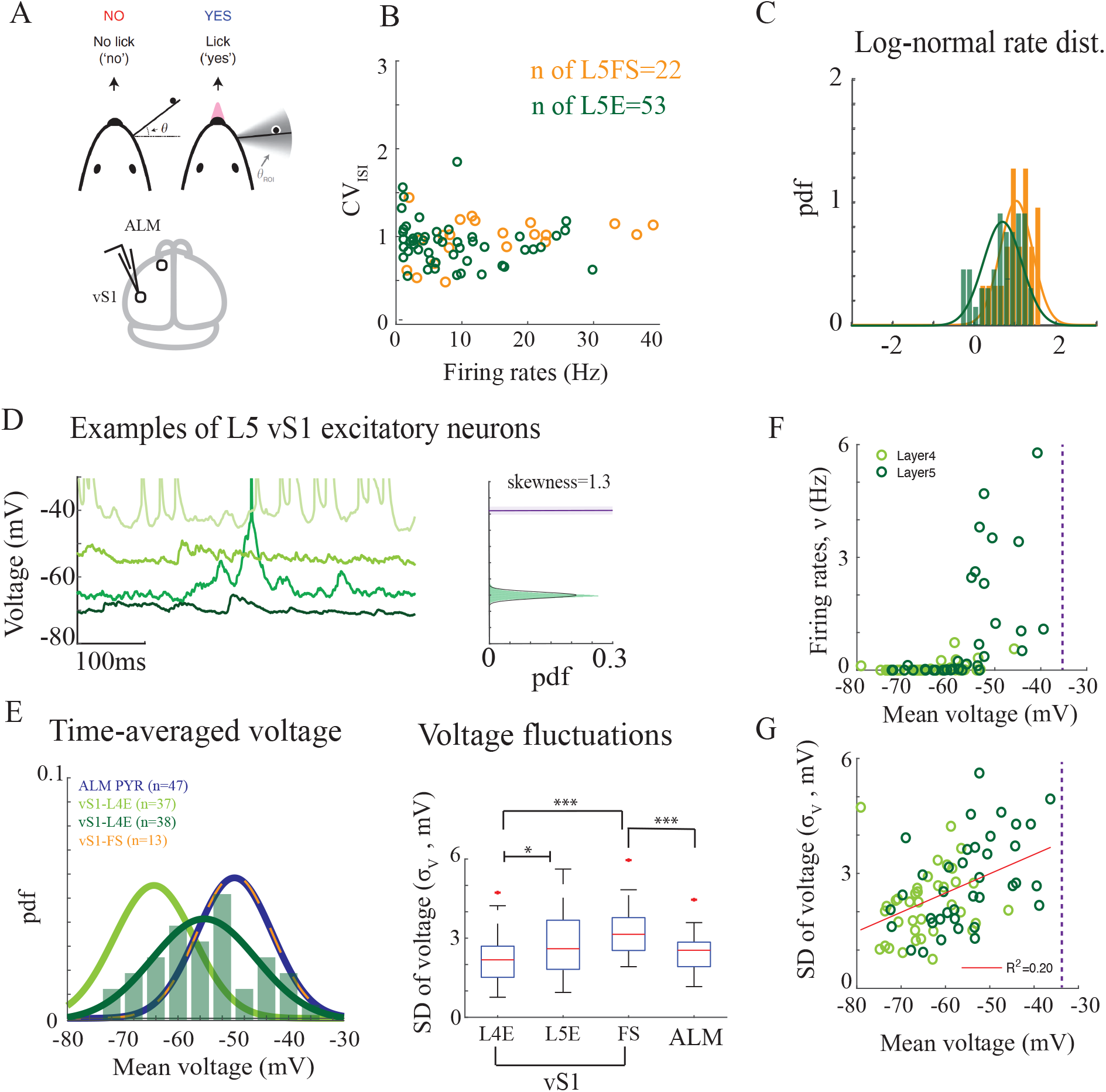
Supra and sub-threshold statistics of layer 5 S1 neurons in mice performing a Go/NoGo task. **A**. Top: behavioral task. Bottom: recording area. **B**. *CV*_*ISI*_ against neuronal firing rates for neurons in layer 5. **C**. Probability density function (pdf) of the log-rates. Solid lines: fit to a Gaussian distribution. **D**. Example of layer 5 excitatory neurons. Left: Activity of example neurons during the first 0.5 second of a whisking episode. Right: Sub-threshold voltage distribution of one of the neurons. Solid line: fit to a Gaussian distribution. **E**. Left: Probability density function (pdf) of time-average voltage for all recorded neurons, including layer 5 excitatory neurons in vS1. Solid lines: fit to a Gaussian distribution. Right: SD of (single neuron) voltage fluctuations for all recorded neurons. **F**. Firing rates vs. mean voltage for the excitatory neurons. **G**. SD of (single neuron) voltage fluctuations against the mean voltage. Red line: linear regression. In (B-C) Extracellular and intracellular recordings. In (D-G) whole-cell recordings. In (B-C,E-F) analysis during non-whisking periods.

**Figure S 2:**
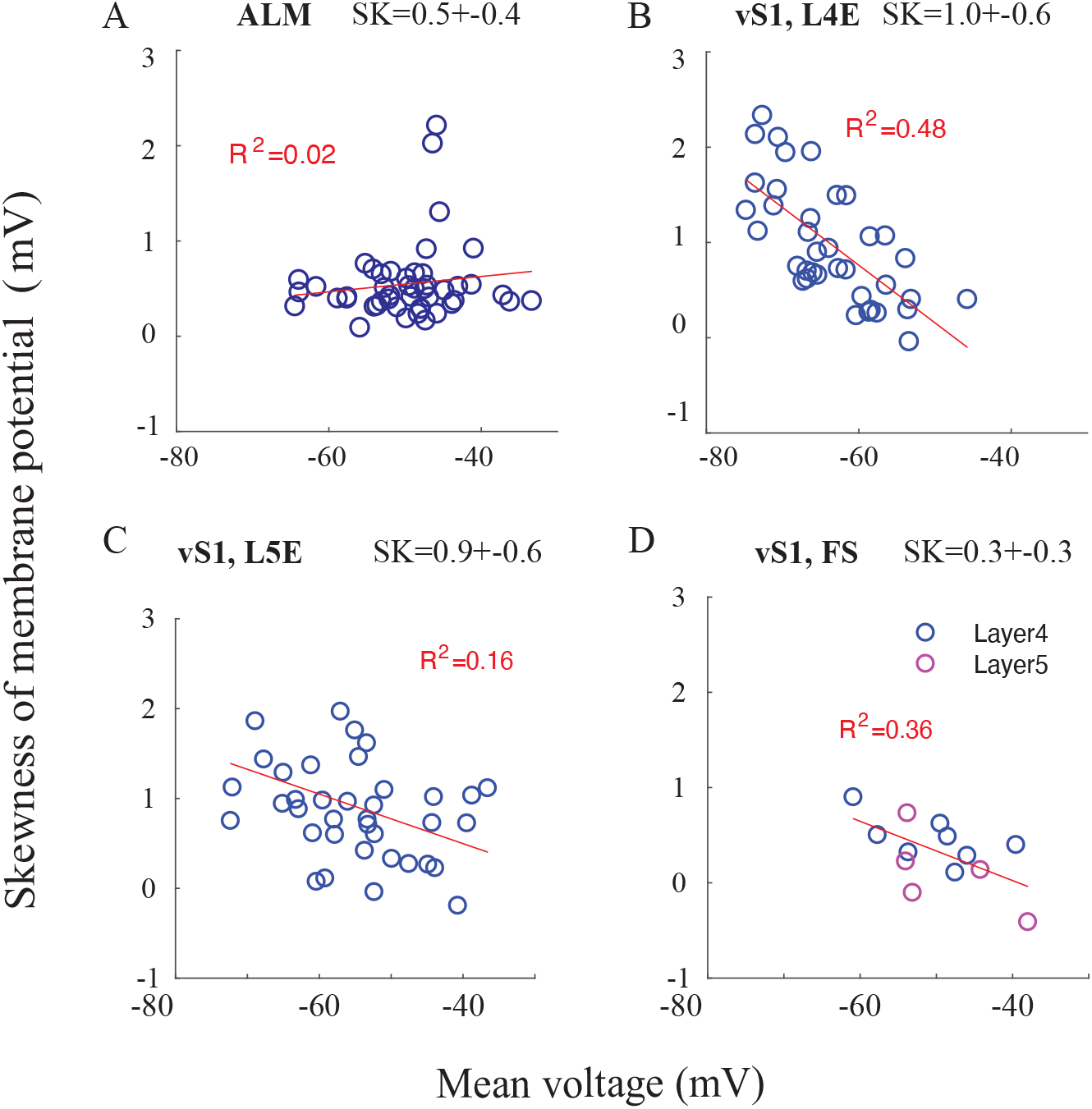
Skewness of membrane potential fluctuations of neurons in ALM and vS1. Skewness of the voltage of a neuron, *V*_i_ (*t*), defined as 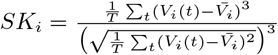, against the mean voltage for ALM (A) and vS1 neurons (B-D). Population-average skewness, 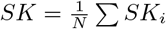, of each population is given in the title.

**Figure S 3:**
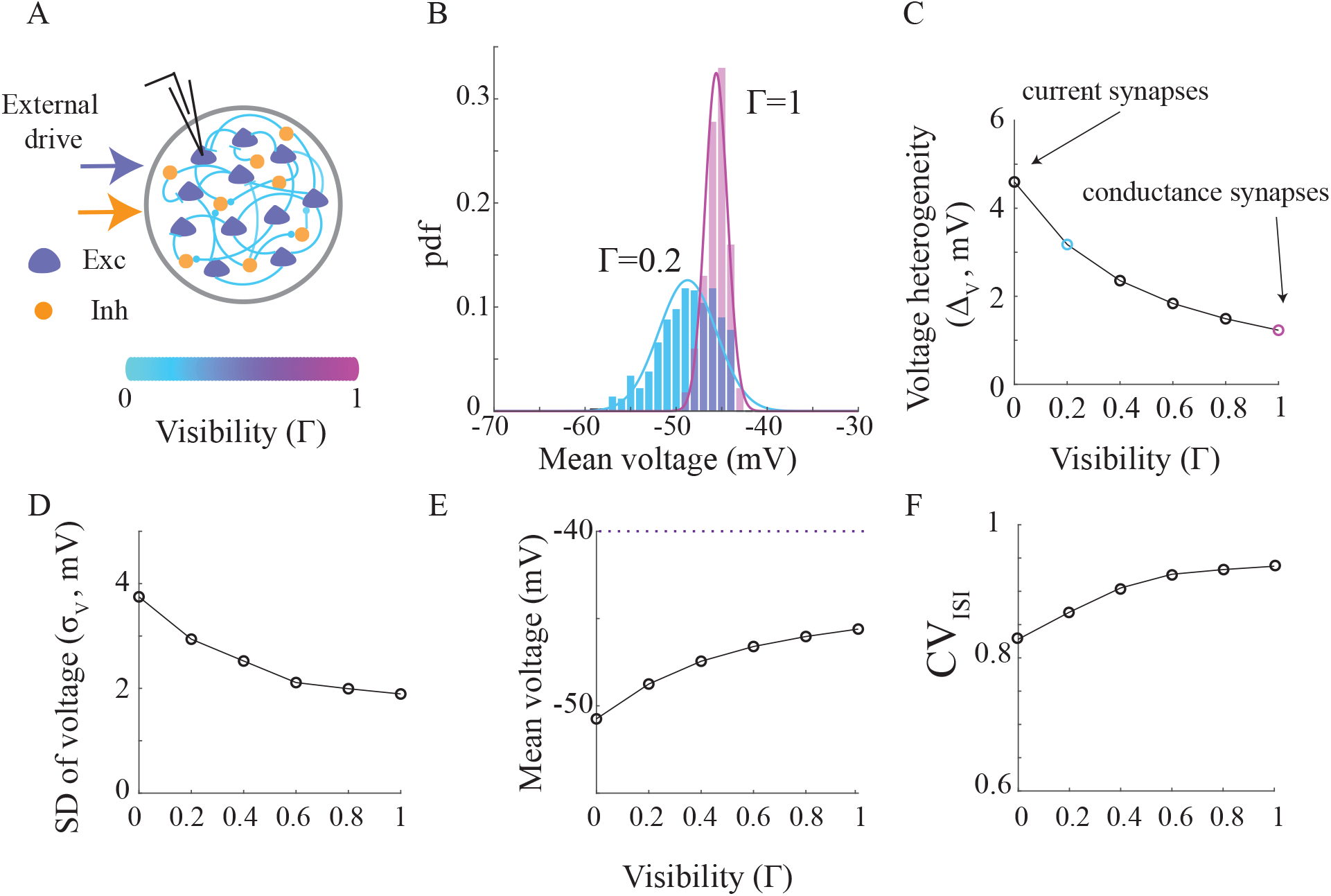
The effect of the visibility of the conductance on sub-threshold heterogeneity and spiking irregularity in a recurrent network. **A**. A cartoon of a network of integrate-and-fire neurons with a mixture of current and conductance synapses. The case in which Γ = 1 corresponds to a pure conductance-based synapse, while the case of Γ = 0 is what is known in the literature as current-based synapses, in which synaptic changes do not affect the conductance of the neuron (see Eq.(12) and Methods). The visibility parameter in this figure is the same for all excitatory, inhibitory and external synapses and we merged the attenuation parameter together with the synaptic strength (*g* in Eq. (12)). **B**. Distribution of time-average voltage of excitatory neurons in a network with Γ = 0.2 (cyan) and of pure conductance-based synapses, Γ = 1 (magenta). Threshold at −40*mV* **C**. Voltage heterogeneity vs. the visibility parameter. Colored circles are the SDs of the distributions of the examples in (B). **D-F**. SD of (single neuron) voltage fluctuations (D), mean voltage (E) and coefficient of variation of the inter-spike-intervals (*CV*_*ISI*_) (F) against the visibility parameter. Note that the SD of the voltage and the *CV*_*ISI*_ varied with Γ in opposite directions. In all panels the mean rate of the excitatory and inhibitory neurons was kept constant (4*Hz* and 9*Hz*, respectively) when changing the visibility parameter by adapting the external inputs.

**Figure S 4:**
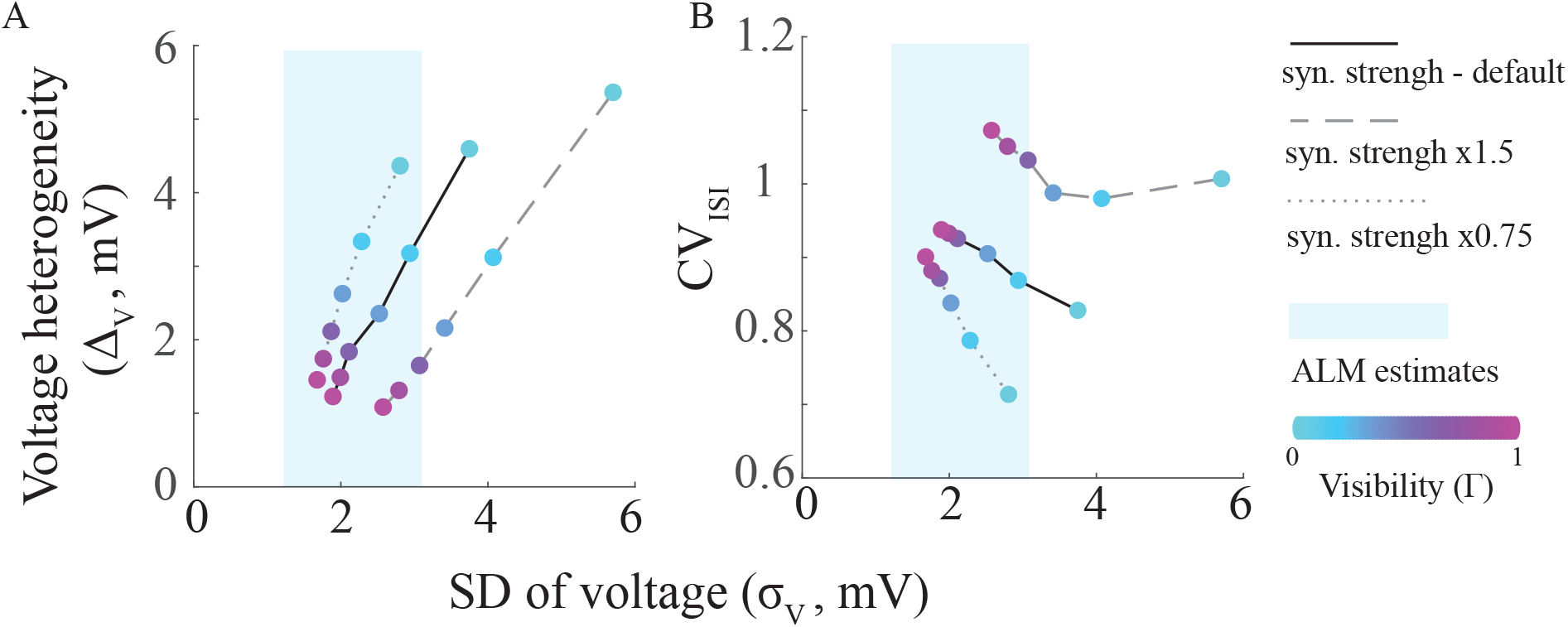
Visibility and network parameters in networks of extended-like point neurons that are consistent with ALM data. Sweep of visibility and network parameters to find visibility parameters that are consistent with ALM data. Solid lines: same figures as in Fig.S3C,D,F, but plotted against the SD of the voltage. The value of each circle is obtained from a network simulation with a different visibility parameter (ranging from zero to one, with jumps of 0.2). Dashed lines: same as solid lines but for a different set of network parameters. To keep the average firing rate in the network constant, and consistent with ALM data, we increased the synaptic strengths by a factor of 1.5 (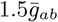 and 1.5*Ī*_*ext,a*_ of the parameters in Table 2). As a result, voltage fluctuations in the network increased, while the mean rate was not changed. Dotted lines: Same as dashed lines but with a 0.75 factor. Cyan: mean±SD of the estimated voltage SD in the population of ALM neurons (see Fig.2E, right). Large voltage heterogeneity is limited by the level of voltage fluctuations and the *CV*_*ISI*_ of neurons in the network. Large voltage heterogeneity together with high *CV*_*ISI*_ in the range of the SD of the voltage of ALM neurons can be achieved for ranges of Γ = 0.2 − 0.4. Note that larger visibility parameters for *σ*_*V*_ ≈ 2*mV* would decrease both the voltage heterogeneity and the *CV*_*ISI*_ to values that are inconsistent with ALM data. **A**. Voltage heterogeneity against voltage SD. **B**. *CV*_*ISI*_ against voltage SD.

**Figure S 5:**
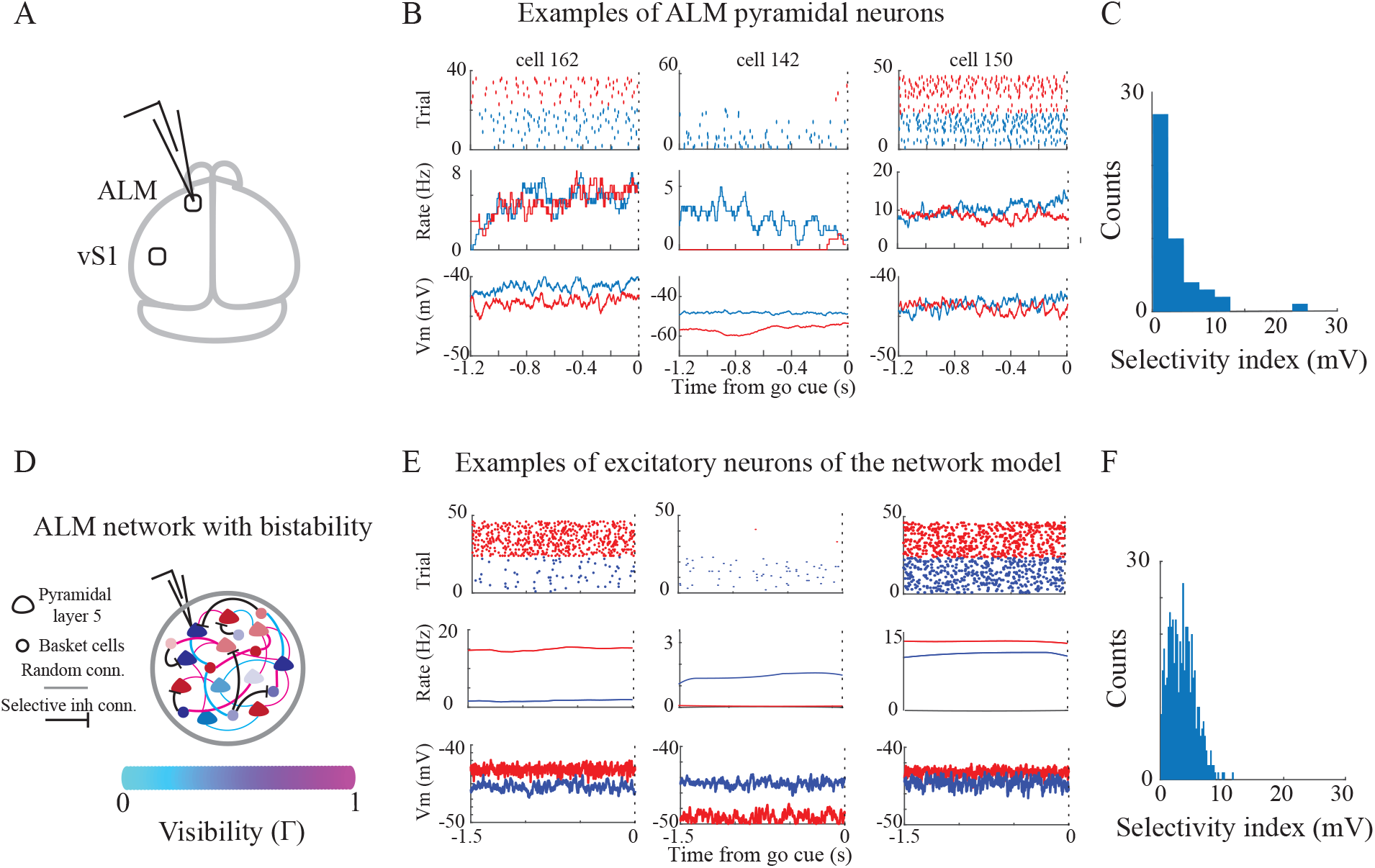
Supra and sub-threshold heterogeneity in selectivity of ALM neurons to licking direction in the experiment and model. ALM data in (A-C) and model in (D-F). **A**. Recording area. **B**. Raster of the spikes (top), peri-stimulus time histogram (PSTH, middle) and average sub-threshold activity (excluding spikes, bottom) for lick right (blue) and left (red) trials in 3 example ALM neurons. PSTHs were calculated using a 10ms window. **C**. Distribution of sub-threshold selectivity to licking direction 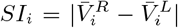, with 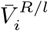 the average voltage for lick left/right of the *i*’th neuron). **D**. Cartoon of a network with extended-like point neurons with visibility and attenuation parameters, estimated from multi-compartment simulations of layer 5 and basket cells. The recurrent connectivity is a combination of a random component and a structured competitive component from the inhibitory neurons that supports bistability in the network. This bistability is the reason for the persistent and selective states in the network (see Methods and [Lebovich et al., 2019]). **E**. Same as (B) but for 3 examples of excitatory neurons in the model. **F**. Same as (C), but for excitatory neurons in the model.

**Figure S 6:**
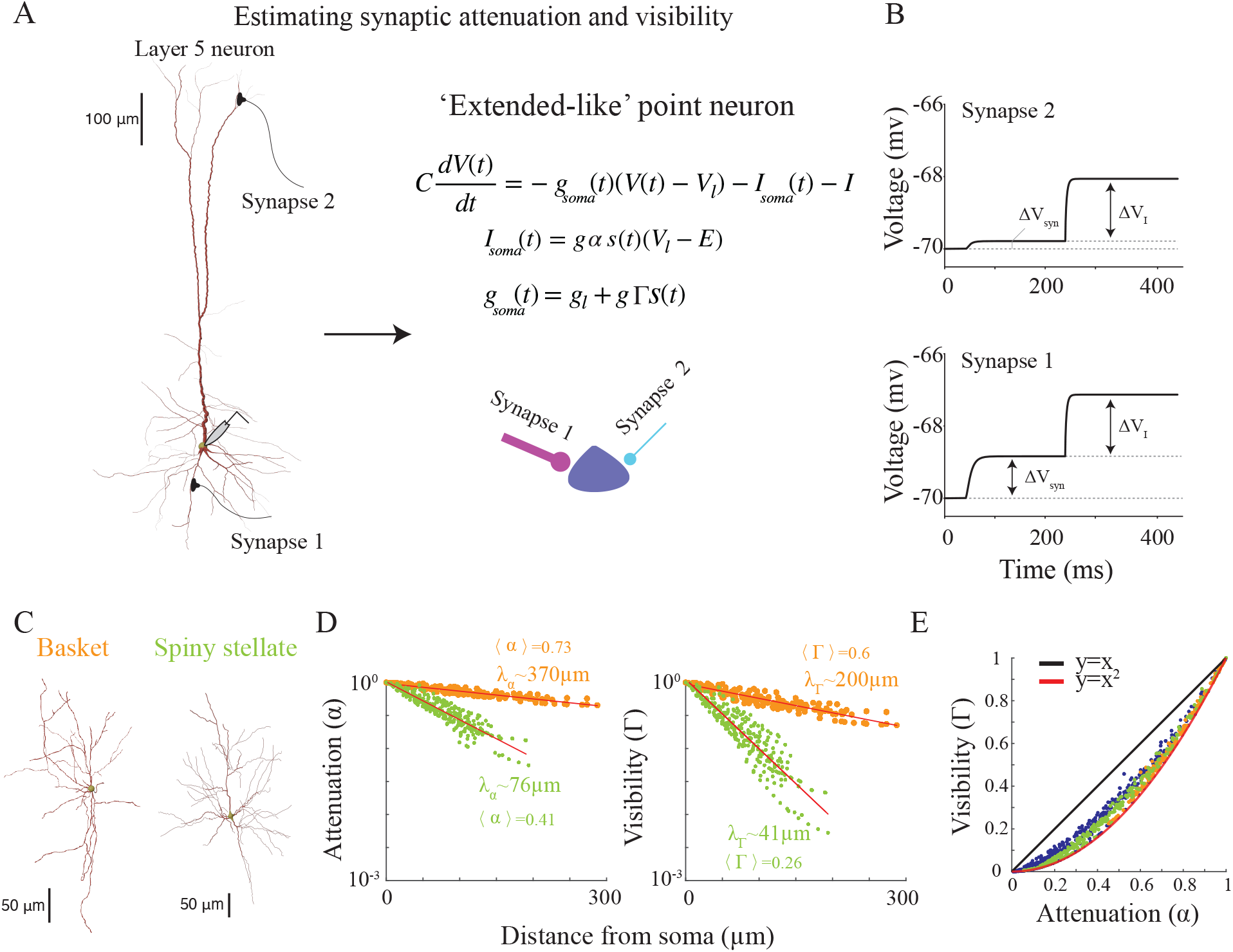
Estimation of the attenuation and visibility for synapses in a multi-compartment model. **A**. Estimation of the attenuation and visibility parameters of the phenomenological point neuron model from a simulation of a multi-compartment model. Left: reconstruction of the simulated layer 5 pyramidal neuron. **B**. The voltage dynamics of the ‘extended-like’ point neuron. Right: The voltage traces of the two example synapses. First, we activate a synapse in a multi-compartment model and measure the change in voltage at the soma (Δ*V*_*syn*_). We then inject a step current at the soma (Δ*I*) and measure the voltage change (Δ*V*_*I*_). The comparison of Δ*V*_*syn*_ and Δ*V*_*I*_ in the multi-compartment model and the point neuron is used to estimate the visibility (Γ) and attenuation (*α*) parameters; see Methods. We repeat this process for every synapse along the dendritic tree. **C**. Same as in (A), but for a multi-compartment model of basket (orange) and spiny stellate (green) cells. **D**. Left: Attenuation vs. the distance of the synapse from soma for the cell in (C). Red line: fit to an exponential decay. Right: Same as left, but for the visibility parameter. Note that the decay rate of the visibility is around twice the decay rate of the attenuation parameter. **E**. The estimated visibility against attenuation for synapses in the three cell types. Note that the visibility always decays faster than the attenuation, and that for an infinite cylinder Γ ≈ *α*^2^ (Fig.S7C; [Koch et al., 1990]).

**Figure S 7:**
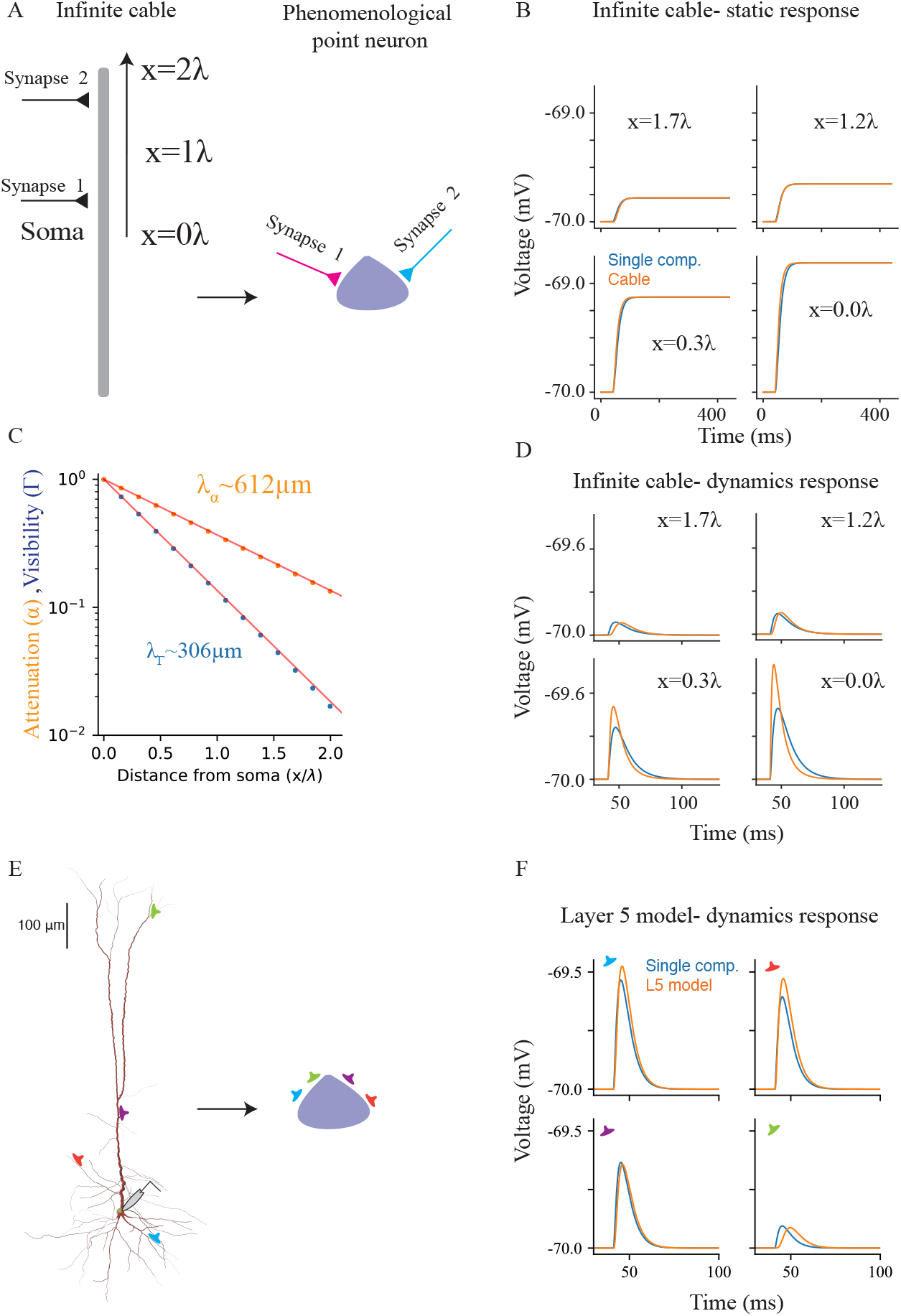
Mapping synaptic activity in an infinite cable and layer 5 neuron to activity of an extended-like point neuron. **A**. Left: a cartoon of the simulated infinite cylinder (with length and diameter of 9500 *μ*m and 1.5*μ*m respectively, and specific membrane resistance of 10,000 Ω*cm*^2^) with a synapse that is located along the cylinder. Right: the extended-like point neuron. **B**. Four examples of the static response of the cable (orange) to activating a synapse at different locations along the cable, and the corresponding mapped synapse when activated on the point neuron (blue). **C**. The attenuation (*α*) and visibility (Γ) of the synapse against the location of the synapse. Note that in an infinite cylinder *α* ∝ *e*^*−x/λ*^ and Γ ∝ *e*^*−x/*2*λ*^ and thus *λ*_*α*_ = 2*λ*_Γ_ [Koch et al., 1990]). **D**. Four examples of the dynamic response of the cable (orange) to activating a synapse (alpha function) at different locations along the cable, and the corresponding mapped synapse of the point neuron (blue). Note that the differences in the voltage are a result of estimating *α*, Γ using the static response of the neurons. **E**. Mapping of the layer 5 neuron (as in main text). **F**. Same as (D), but for the layer 5 neuron in (E).

**Figure S 8:**
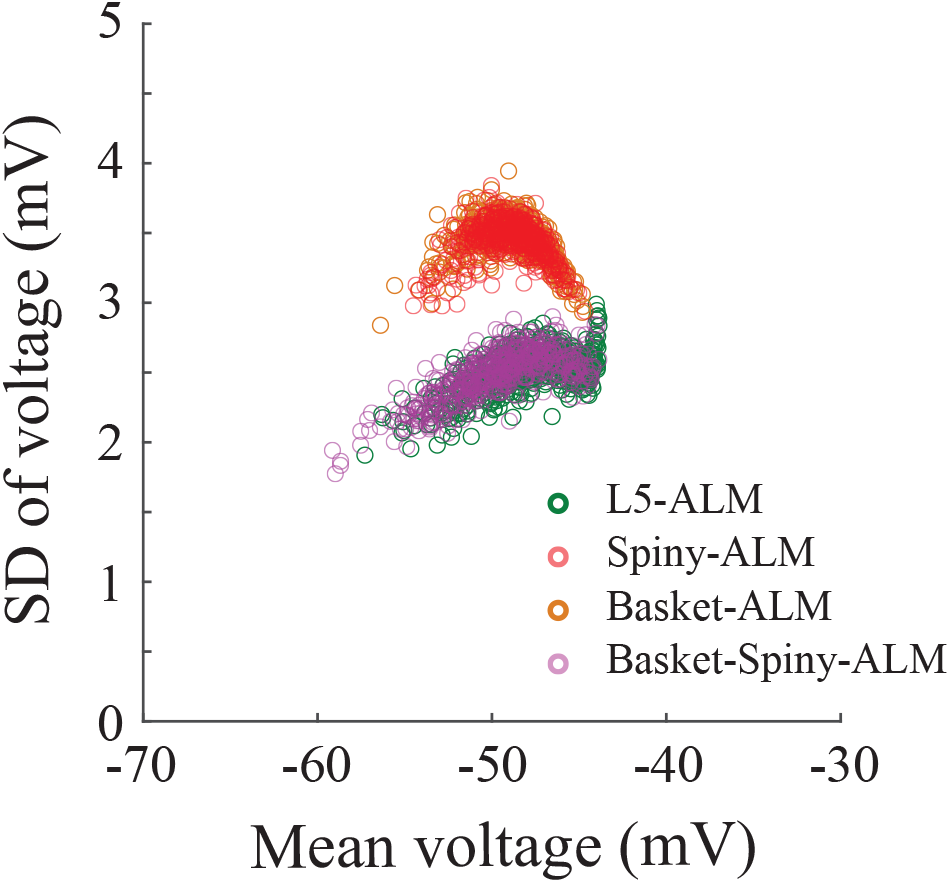
Voltage statistics of neurons in a network of extended-like neurons is similar when the visibility and attenuation parameters are estimated from a layer 5 pyramidal neuron or from a spiny stellate neuron. SD of voltage fluctuations against the mean voltage of 500 neurons in a network with layer 5-basket cells (green and orange) and spiny stellate-basket cells (purple and red). Same network parameters as in Fig.5A-E.

**Figure S 9:**
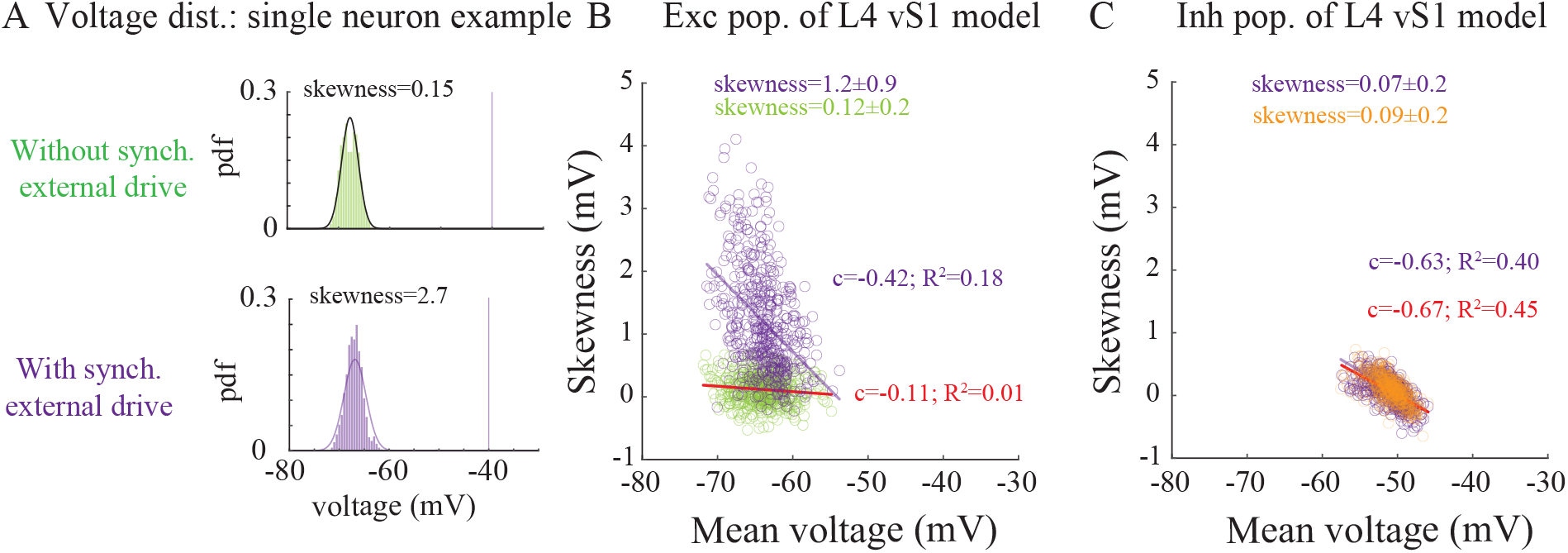
Deviations from Gaussianity of voltage fluctuations in a model of layer 4 vS1 network. **A**. Example of an excitatory neuron in layer 4 vS1 network model without (top-green) and with (bottom-purple) synchronous external drive (see Methods). Without external drive the voltage distribution is well-fitted with a Gaussian distribution (solid line; low skewness). **B**. Skewness of excitatory neurons without (green) and with (purple) synchronous external drive. Red line: linear fit in a network without synchronous external drive. Correlations between skewness and mean voltage of neurons in the network are low (small c). Purple line: linear fit in a network with synchronous external drive. Skewness and mean voltage of neurons in the model is negatively correlated (negative c). Note that the synchronous external drive mainly affects the hyperpolarized excitatory neurons. **C**. Same as (B), but for the inhibitory population, which are more depolarized than the excitatory neurons. The external drive only weakly affects the skewness of the neurons.

**Table 2:**
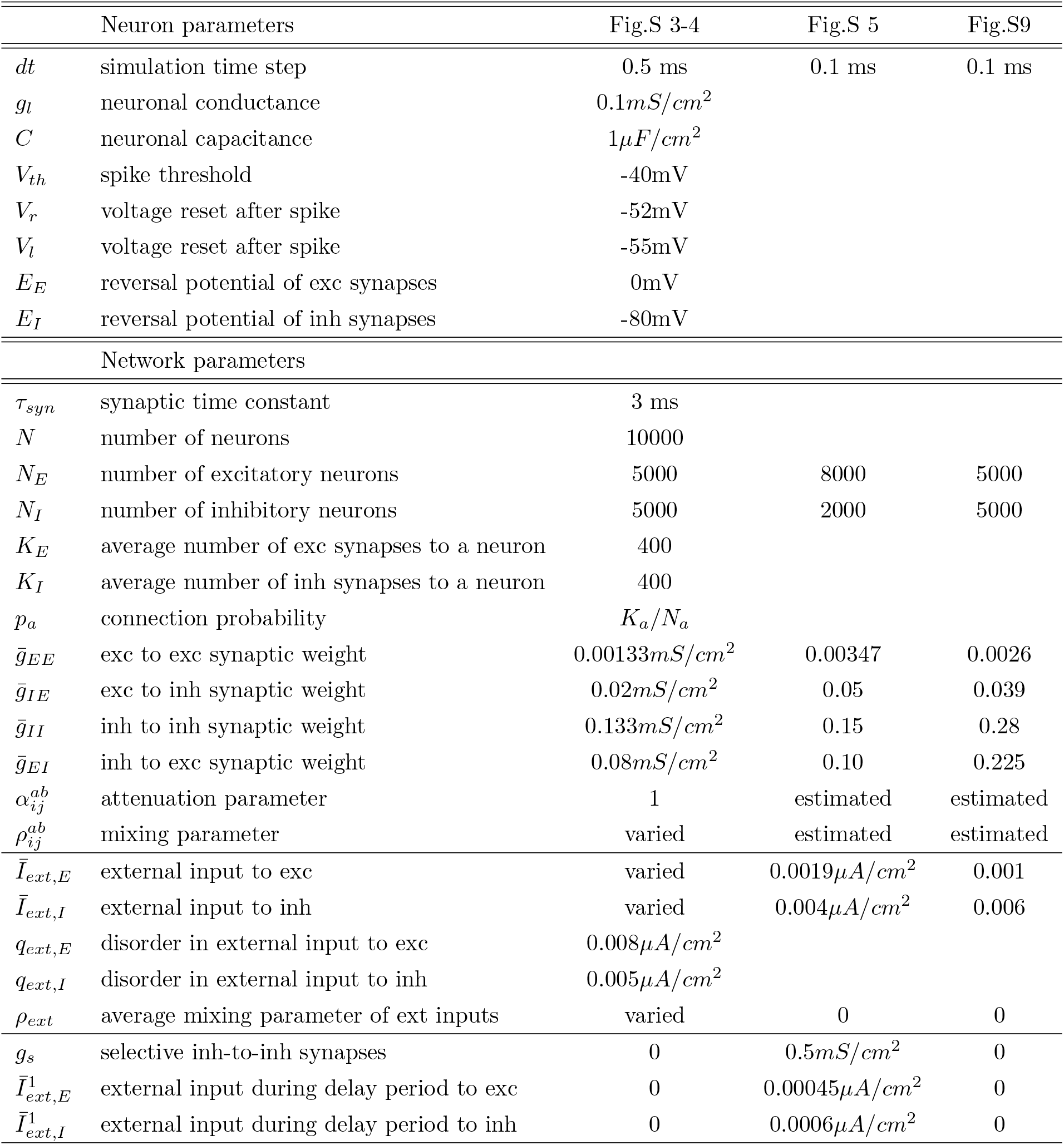
Network simulation parameters for supplementary figures.e

## Supplementary materials

### Network architecture for a balanced network that supports persistent activity through bistability

Neurons in ALM tend to exhibit persistent, or slow, dynamics during the delay period [Inagaki et al., 2019]. To model this, we add to the random connectivity of ALM network model additional structured connections as in [Lebovich et al., 2019]. These additional structured connections allowed the balanced network to be bistable without breaking the EI balance.

We follow [Lebovich et al., 2019] and model an additional selective feedforward input into each population, excitatory or inhibitory, that consists of two subsets of neurons, namely left (L) and right (R) selective neurons. Before the instruction appears this additional feedforward input, 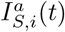, is zero for all neurons. Upon presentation of a right stimulus, 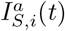, into R-selective neurons is stronger than for the L-selective neurons, and vice versa, while during delay period this input is constant, and equal, to both L and R populations. Specifically, instead of Eq.(15) we take

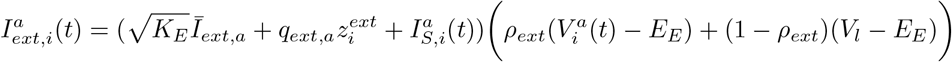

with the input

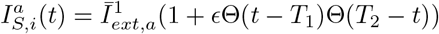

that is non-zero only during sample and delay periods (that last 2100*ms*). The parameter *c* is positive for R trials and negative for L trials, Θ(*x*) is the Heaviside function, *T*_2_ − *T*_1_ is the sample duration (100*ms* in simulations) and with 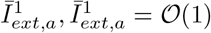. Thus, there is no selective feedforward input to neurons in the model during delay period.

We denote the subset of R-selective (L-selective) neurons in population *a* by *R*^*a*^ (*L*^*a*^). Neuron (*i, a*) ∈ *R*^*a*^ if 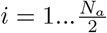 and (*i, a*) ∈ *L*^*a*^ if 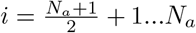.

The recurrent connectivity in this extended model has two components. One is functionally specific and the other is not. The non-specific component is fully random, Erdős–Rényi (ER) graph, and does not depend on the selectivity of the pre- and post-synaptic neurons. The competition between the left (L) and the right (R) selective neurons is mediated by an additional set of connections. These connections are specific and are much less numerous, but stronger than the unspecific ones. There are no specific connections from the excitatory neurons to other neurons. The probability for a specific connection from an inhibitory neuron to another excitatory or inhibitory neurons is 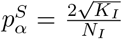. Therefore, each neuron (excitatory as well as inhibitory) receives, on average, 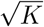 connections from inhibitory neurons whose selectivities are different from its own, and on average *K* non-selective inhibitory connections.

The strength of the specific connections depends solely on the neurons’ type, *g*_*aI*_*g*_*S*_, with *g*_*S*_ determining the strength of the selective synapses with respect to the non-selective, random, connections. It is non-zero only in the simulations of Fig.S5. The total current into neuron (*i, α*) due to the recurrent interactions follows Eq.(14), with the recurrent connections given by 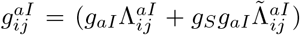, with 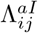 the adjacency matrix of an ER graph with 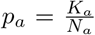, and 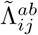 the adjacency matrix of another ER graph with 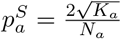.

With this architecture and feedforward input both the excitatory and inhibitory neurons in the network are selective to the licking directions, both during sample, and without selective feedforward input during delay period.

### Mean-field analysis of sub-threshold statistics of conductance-based integrate-and-fire neurons

Here we consider the case of a conductance-based integrate-and-fire neuron and calculate the distribution of mean and voltage SD. By inserting Eqs (23) into (13), one can use a Fokker-Plank formalism to analytically describe the firing rate and the membrane potential distribution of a conductance-based neuron. We follow here [Richardson, 2004], and add the neuronal index that arise due to the heterogeneity in synaptic inputs. It is useful to first introduce several quantities, where to simplify notations we write them without their postsynaptic population index:

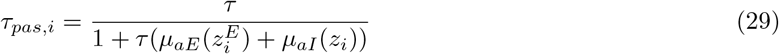

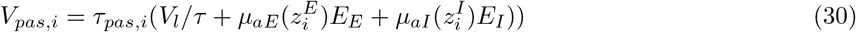

that only weakly depend on the disorder in the network because the disorder is always an order 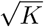 smaller than the mean *μ*_*ab*_. In fact, in the large *K* limit Eq.(29) goes to zero and Eq.(30) is independent of *K*.

Furthermore, we also introduce two voltage quantities:

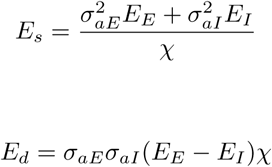

and

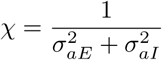

which none of them depends on the quenched disorder. Finally, we also introduce the dimensionless parameter 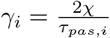, which needs to be large in order for the Gaussianity assumption to hold.

Using a linear change of variables from voltage to a dimensionless variable, *x*_*i*_:

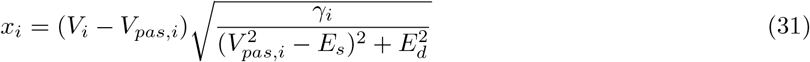

with *x*_*th*_ and *x*_*r*_ corresponding to placing *V*_*th*_ and *V*_*r*_ in Eq.(31). It is then possible to write the equilibrium distribution of the FP equation for *f* (*x*_*i*_):

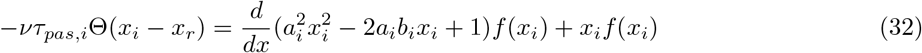

with 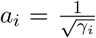 and 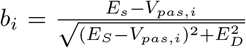. Integrating Eq.(32) after multiplying it with 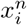 gives, after notorious algebra, the moments of the distribution. The first and second moments of the voltage yields:

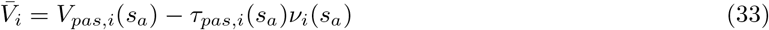

and

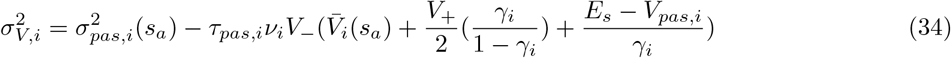

with the variance of a passive conductance-based IF neuron given by 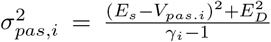. One should note that as *γ*_*i*_ scales with the synaptic strength 1*/g*_*ab*_, we get that when *g*_*ab*_ is small, and for which the Gaussianity assumption is valid, the leading order of the fluctuations takes the simple form

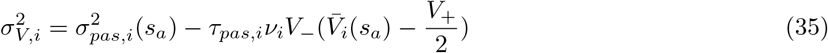

Thus, both the mean and the SD of the voltage for conductance-based IF neurons (Eq.(33),(35)) have the same form as the mean and SD of the current-based IF neurons. It is the mean or SD of the passive neuron, together with another term that is proportional to the rate of the neuron.

